# Differentiating Drosophila female germ cells initiate Polycomb silencing by altering PRC2 sampling

**DOI:** 10.1101/2020.02.03.932582

**Authors:** Steven Z DeLuca, Megha Ghildiyal, Wanbao Niu, Liang-Yu Pang, Allan C. Spradling

## Abstract

Polycomb silencing represses gene expression and provides a molecular memory of chromatin state that is essential for animal development. We show that *Drosophila* female germline stem cells (GSCs) provide a powerful system for studying Polycomb silencing and how it is established. GSCs resemble pluripotent mammalian embryonic cells in lacking silenced chromatin, but most GSC daughters, like typical somatic cells, induce Polycomb silencing as they differentiate into nurse cells. Developmentally controlled changes in the levels of two Polycomb repressive complex 2 (PRC2)-interacting proteins, Pcl and Scm, initiate differentiation. In germline stem cells, abundant Pcl inhibits silencing by slowing PRC2 and diverting it from PRE sequences. During differentiation, core PRC2 represses inactive loci while Scm and residual Pcl cooperate to enrich PRC2 and silence traditional Polycomb domains. We propose that PRC2-interacting proteins regulate the transition from a variable to stable transcription state during differentiation by altering the rate that PRC2 samples regulatory sequences.

## Introduction

Differentiation is the defining mechanism enabling the evolution and development of multicellular animals. During early *Drosophila* embryonic development, cascades of transcription factors transform two initial body axes into a precise coordinate system that identifies nearly every cell by a unique combination of factors based on their position (Fowlkes et al., 2008; Karaiskos et al., 2017; St Johnston and Nüsslein-Volhard, 1992). Further elaboration of a differentiation program requires the acquisition of a cellular “memory” mediated by an exceptional form of repression known as Polycomb silencing (Jones and Gelbart, 1990; Struhl and Akam, 1985; Wedeen et al., 1986). Initially characterized by genetic studies of Hox gene regulation along the anterior-posterior axis of the *Drosophila* embryo (Lewis, 1978), Polycomb group gene (PcG-gene) products recognize repressed loci, coat kilobases of repressed enhancer regions (PcG domains), limit subsequent transcription, and restrict eventual cell fates (Schuettengruber et al., 2017). Subsequent research revealed that Polycomb silencing is also utilized by mammalian embryos and likely by all animals, and programs the differentiation of all somatic embryonic cells as well as progeny cells downstream from pleuripotent embryonic stem cells (ESCs) (Aloia et al., 2013; Montgomery et al., 2005). Despite these critical roles, the mechanisms PcG proteins use to initially recognize target sites, induce silencing, and maintain a memory of silencing in descendent cells is imperfectly understood. In particular, learning how PcG proteins interact with their targets in PcG domains at the time cellular “memories” are initially formed is critically important to addressing these questions.

PcG genes encode two major protein complexes, Polycomb Repressive Complex 1 (PRC1), an E3 ligase that ubiquitylates histone H2A on lysine 119 (H2AK119ub) (Franke et al., 1992; Shao et al., 1999; Wang et al., 2004a), and PRC2, which methylates histone H3 on lysine 27 (H3K27me1/2/3) (Cao et al., 2002; Czermin et al., 2002; Kuzmichev et al., 2002; Müller et al., 2002). In fly embryos, silencing and PRC2 and H3K27me3 enrichment on PcG domains first appears as anterior-posterior patterning is being completed during a massive upregulation of zygotic gene expression during cell cycle 14 (Alhaj Abed et al., 2018; Li et al., 2014; Pelegri and Lehmann, 1994). In preimplantation mouse embryos, H3K27me3 is distributed in a “non-canonical” pattern throughout gene deserts and inactive loci (Liu et al., 2016; Zheng et al., 2016) and is not further enriched on its canonical sites: CpG-rich “islands” (CGIs) around enhancers and promoters of developmentally-induced genes (Ku et al., 2008; Tanay et al., 2007). However, H3K27me3 is highly enriched on CGIs in mouse embryonic stem cell cultures derived from preimplantation embryos (Boyer et al., 2006). The transition from the non-canonical to canonical H3K27me3 pattern likely involves the initial PRC2 specificity-establishing events, but this transition remains poorly understood. Unfortunately, studying the establishment of PRC2 specificity in early embryos is hampered by difficulties in purifying and genetically manipulating a short-lived transition state controlled by both a maternal and zygotic supply of PcG proteins. Nevertheless, many studies have investigated factors controlling PcG targeting specificity in other cell types in both flies and mammals, which may offer clues into how PcG targeting specificity is first induced in early embryos (recently reviewed in (Kassis et al., 2017; Kuroda et al., 2020; Laugesen et al., 2019; Yu et al., 2019).

One prominent difference between PcG silencing in flies and mammals are the sites within target genes that appear to recruit PcG proteins. In flies, PhoRC binds specific sites in Polycomb Response Elements (PREs) and recruits PRC1 and PRC2 to PcG domains (Brown et al., 1998; Busturia et al., 2001; Fritsch et al., 1999; Klymenko et al., 2006; Mishra et al., 2001; Shimell et al., 2000). PREs are enriched in several hundred broad H3K27me3-coated loci including the Hox clusters and the enhancers of other developmentally regulated transcription factors and signaling components (Schwartz et al., 2006). Mammals, however, lack a prominent PhoRC-like targeting mechanism and instead enrich PcG proteins on CGIs to silence a larger number of tissue-specific genes than flies (Jermann et al., 2014; Ku et al., 2008; Lynch et al., 2012; Mendenhall et al., 2010; Tanay et al., 2007). Relatively few of these mammalian PRC2-regulated genes are clustered like the four Hox loci.

In mammals, multiple mechanisms have been proposed to contribute to PRC1 and PRC2 recruitment to CGIs including DNA shape, nucleosome spacing, histone tail modifications including a “bivalent” combination of H3K27me3 and H3K4me3, and interactions with RNA (Yu et al., 2019). Notably, mouse Kdm2 subunits of PRC1 and the Pcl subunits of PRC2 specifically bind CGI sequences *in vitro* and have been proposed to initially recruit each complex to CGIs *in vivo* (Farcas et al., 2012; He et al., 2013; Li et al., 2017; Perino et al., 2018; Wu et al., 2013).

Additionally, Pc proteins in PRC1 complexes bind the H3K27me3 catalyzed by PRC2 while the Jarid2 accessory subunit of PRC2 binds H2AK119ub catalyzed by PRC1 (Cao et al., 2002; Cooper et al., 2016; Czermin et al., 2002; Müller et al., 2002). Because both PRC1 and PRC2 can directly interact with DNA and the histone modifications catalyzed by the other complex, each complex could putatively recruit the other or synergize to promote higher recruitment at overlapping sites. Flies have homologues of Pcl and Kdm2 proteins but fly Pcl lacks sequence specificity *in vitro* (Choi et al., 2017) and fly Kdm2 is dispensable for viability (Shalaby et al., 2017; Zheng et al., 2014). Furthermore, while PRC1-catalyzed H2AK119ub enriches PcG proteins in mouse and fly embryonic cell cultures, (Blackledge et al., 2014; 2019; Cooper et al., 2014; Fursova et al., 2019; Kahn et al., 2016), it has a minimal effect on PRC2 localization and Polycomb silencing in many fly tissues (Gutiérrez et al., 2012; Pengelly et al., 2015). Instead, disruption of PhoRC or Scm, which links PhoRC to both PRC1 and PRC2 (Frey et al., 2016; Kim et al., 2005; Klymenko et al., 2006; Peterson et al., 2004), causes stronger PcG protein enrichment defects in flies (Brown et al., 2018; Kang et al., 2015; Wang et al., 2004b).

However, it remains possible that some differences between mammalian and *Drosophila* PcG targeting are not intrinsic, but arise from differences in the experimental systems used for their study. For example, scml1, one of four mouse Scm paralogues, promotes H3K27me3 enrichment on CGIs in the male germline, raising the possibility the Scm is a conserved PRC2 targeting factor (Maezawa et al., 2018). Previously, *Drosophila* Pcl was found in a small fraction of embryonic PRC2 complexes, and Pcl was proposed to promote PRC2 and H3K27me3 enrichment near PREs (Nekrasov et al., 2007; O’Connell et al., 2001; Savla et al., 2008; Tie et al., 2003). In mouse embryonic stem cells, Pcl orthologues similarly promote H3K27me3 on CGIs, and the winged helix and tudor domains of mouse Pcls have been proposed to interact with DNA and methylated histones to target Pcl-PRC2 complexes to particular CGIs (Li et al., 2017; Perino et al., 2018).

We have established a new system, *Drosophila* female germ cell differentiation, for analyzing Polycomb silencing (Fig. 1A) that avoids the cellular and genetic complexity of early embryonic development. Female germ cells in Drosophila, mouse and diverse other species (Lei and Spradling, 2016; Matova and Cooley, 2001), not only give rise to oocytes, but mostly produce a late-differentiating cell type known as nurse cells that nourish the oocytes by donating cytoplasmic organelles, RNAs and proteins before undergoing programmed cell death. In Drosophila, nurse cells differentiate downstream from germline stem cells (GSCs) and undergo polyploidization, which makes them abundant and easy to purify for genomic studies at every step in the differentiation process. Whether nurse cells are true somatic cells that differentiate using canonical PcG silencing is controversial because germline mutation of some PcG-group genes (*E(z)* and *Su(z)12*) disrupt nurse cell – oocyte differentiation while mutation of most others do not (Iovino et al., 2013).

**Figure 1:**
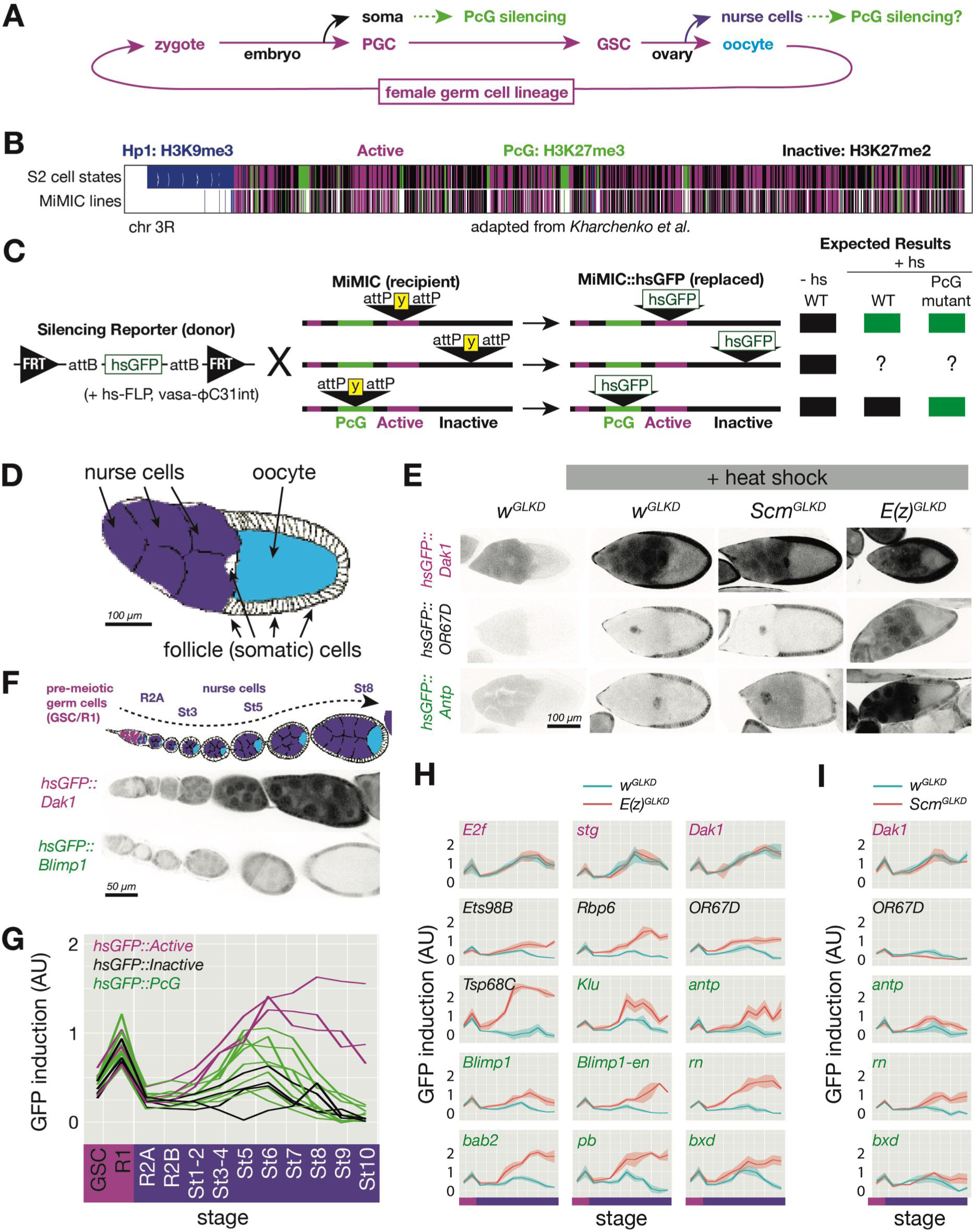
Developmentally regulated silencing in inactive and PcG domains in the germline A) Cyclical lineage of female germ cells and two dead-end derivative lineages, soma and nurse cells. (B) Map of chromosome 3R showing color-coded, simplified chromatin states. (C) Integration protocol of hsGFP silencing reporters into MiMIC insertions within different chromatin domains and expected expression +/-heat shock (hs). (D) Cell types in a stage 10 follicle. GLKD should affect nurse cells (purple), and oocytes (cyan) but not surrounding somatic cells (white). (E) Stage 9/10 follicles showing GFP fluorescence from reporters integrated in active (near *Dak1*), inactive (near *OR67D*) or PcG (near *Antp*) chromatin, in germline knockdown (GLKD) of control (*w*), *Scm,* or *E(z)*. Somatic follicle cells serve as an internal control. (F) Diagram of germline development from pre-meiotic stages (GSC/R1, pink). Nurse cells (purple) differentiate from oocytes (cyan) in region 2A (R2A); nurse cells and oocytes grow further (St3-St8). Below, GFP fluorescence after heat shock from two indicated lines. (G) Plot of mean GFP induction ([GFP]^+hs^ – [GFP]^-hs^) in nurse cells or their precursors across 12 developmental stages for 15 reporter lines colored according to their chromatin domain. (H-I) The effect of *E(z)^GLKD^* (H) or *Scm^GLKD^* (I) on reporters near the indicated genes colored by domain type. Solid line indicates mean fluorescence; shading shows 1 standard deviation from the mean. X-axes colored for stage as in G. Size bars: D,E 100μ; F 50μ.

Here, we show that nurse cells acquire canonical Polycomb silencing during differentiation using similar mechanisms as early embryos, making female germ cells a powerful system for studying Polycomb silencing. Nurse cell progenitors lack silencing and contain a similar “non-canonical” H3K27me3 pattern as early embryos. Complete silencing of PcG domains in differentiated nurse cells requires multiple PcG proteins, including components of PRC1, in addition to core subunits of PRC2. PcG gene mutations are less disruptive in germ cells compared to embryonic cells, because interfering with the single, relatively simple nurse cell program affects oocyte completion more weakly than disrupting myriad, interdependent somatic cell type differentiation programs in a developing embryo. Finally, we show how two developmentally regulated PcG proteins alter PRC2 targeting to initiate silencing during differentiation. Our results suggest a specific model for the establishment of Polycomb silencing in naïve precursors, and provide new insights into how Polycomb silencing has evolved to maintain its conserved function in animals such as mammals and insects.

## Results

### A system of reporters to analyze developmental gene silencing

To examine the suitability of Drosophila germ cell differentiation for studies of Polycomb silencing initiation, we first developed a method to detect and measure silencing throughout the genome in single cells. Because removing silencing may not robustly induce endogenous gene transcription in the absence of transcriptional activators, reporter-based approaches have been used to document repression within PcG-silenced and Hp1-silenced chromatin *in vivo* (Babenko et al., 2010; Fitzgerald and Bender, 2001; Wallrath and Elgin, 1995; Yan et al., 2002). However, existing reporters were not suitable for probing repressive domains in germ cells for a number of technical reasons. Therefore, we developed a new reporter compatible with female germ cells and easily targeted to potentially silenced loci. Our reporter (hsGFP) consists of a minimal fragment of the Hsp70A gene containing a heat shock-inducible enhancer, promoter, and short 5’UTR fused to Green Fluorescent Protein (GFP) and a transcriptional terminator (Fig1C). We chose the heat shock enhancer and promoter because of its low basal activity, robust inducibility in nearly all cells types, and similarity to promoters of developmentally activated genes (Guertin et al., 2010; Muse et al., 2007; Zeitlinger et al., 2007). Importantly, our reporter was insensitive to ovarian small RNAs derived from *Hsp70* loci due to a truncated 5’UTR (DeLuca and Spradling, 2018).

We used a previously described crossing strategy employing both FLP-mediated and phiC31-catalyzed site-specific recombination (Nagarkar-Jaiswal et al., 2015) to efficiently introduce the silencing reporter into 109 genomic sites that already contained a single MiMIC transposon located within chromatin predicted to be active or repressed (Fig. 1B,C). To identify potentially silenced regions throughout the *Drosophila* genome, we simplified a previous chromatin annotation (Kharchenko et al., 2011) into 3 types of potentially silenced domains– an H3K27me3-enriched “PcG” type (Fig1B green), an H3K9me3-enriched “Hp1” type (Fig1B blue), and an H3K27me2-enriched generally “inactive” type (Fig1B black), and a single type of “active” domain depleted for H3K9 or K27 methylation (Fig1B magenta). From thousands of potential reporter integration sites (Fig1B), we focus here on data from 3 active-, 6 inactive-, and 13 PcG-domain-localized inserts. Data from Hp1-silenced reporters will be reported elsewhere.

Our reporter enables simple tests for PcG-repressed chromatin at specific reporter insertion sites, with single cell resolution, and at nearly every stage of development. While the very different sizes and metabolic activities of different cell types may influence the amount of GFP produced from reporters, our system should reliably detect PcG silencing as long as comparisons are made in a single cell type between reporters in PcG versus non-PcG domains, or between *control* versus *PcG mutant* genotypes. For example, a reporter integrated into a repressed PcG domain should be less inducible than one integrated into an active domain, and genetically removing PcG proteins should increase the induction of reporters in repressed PcG domains but not active domains (Fig1C).

When applied to ovarian germline cells, the reporter system fulfilled these expectations. We compared hsGFP induction in nurse cells at stage 9-10 (Fig. 1D) for an insert in an active (*Dak1*), PcG (*Antp*), or inactive (*OR67D*) region (Fig. 1E). The reporter integrated near *Dak1* was strongly induced in nurse cells following heat shock, while the reporters integrated near *Antp* or *OR67D* did not induce following heat shock. Finally, **G**erm**L**ine-specific RNAi **K**nock **D**own (GLKD) of *E(z)* (*E(z)^GLKD^*) largely relieved repression of the reporters near *Antp* and *OR67D*, and *Scm^GLKD^* partially relieved reporter repression near *Antp* (Fig. 1E). These results suggest that E(z)-dependent silencing impacts not only PcG domains but also inactive domains whereas Scm contributes to PcG but not inactive domain silencing. We extended our analysis to measure reporter induction in 12 additional PcG domains and 2 additional control active domains. We noted that in all cases, reporters in active domains were highly induced while reporters in PcG domains were nearly uninducible in stage 9-10 nurse cells (Fig1G).

### PcG silencing and nurse cell differentiation

To address when Polycomb repression arises during nurse cell development, we measured GFP induction in reporters in active, inactive, and PcG domains throughout germline development, including two stages before (Fig 1F-I, pink bar on x-axis) and 10 stages during and after nurse cell differentiation (Fig1F-I, purple bar on x-axis). We observed very little difference in reporter induction between active and repressed loci in germ cells prior to the onset of meiosis in region 2A (Fig.1G). However, we detected significant reporter silencing in all inactive and some PcG domains by stage 1-2 (Fig1G). We additionally combined reporters with *E(z)^GLKD^* to detect E(z)-dependent silencing with high sensitivity (Fig. 1H). While reporters in active chromatin were not affected by *E(z)^GLKD^* at any stage, we reliably detected reporter silencing in all inactive and most PcG domains by stage1-2, and in all PcG domains by stage 6. E(z)-dependent reporter repression appeared to strengthen in all inactive and PcG inserts as nurse cell development progressed (Fig. 1H). We conclude that PRC2-dependent reporter gene silencing is absent in germline progenitors, induced in nurse cells after they differentiate from progenitors and oocytes, and strengthened as nurse cells mature.

### PRC2 suppresses PcG domains and inactive chromatin domains by distinct mechanisms

How could PRC2 repress reporters outside of PcG domains, which lack high levels of H3K27me3 and PRC1 enrichment? Inactive domains are enriched for PRC2-catalyzed H3K27me2 nucleosomes (Kharchenko et al., 2011), which were previously proposed to antagonize transcription by preventing H3K27 acetylation (Ferrari et al., 2014; Lee et al., 2015). Thus, PRC2 may provide two repressive functions, an H3K27me1/2-dependent function that simply opposes acetylation and an H3K27me3-dependent function that additionally recruits PRC1 and enhances silencing. We hypothesize that the H3K27me1/2-dependent function would act at all repressed domains, while the H3K27me3-dependent function would only act within PcG domains.

To separate H3K27me3- from H3K27me1/2-mediated effects, we screened an RNAi collection targeting known PcG proteins in order to identify genes specifically promoting H3K27me3 on PcG domains (Fig. 2A). In whole mount wild type nurse cells, H3K27me3 appeared as intense puncta that stand out from a hazy general chromatin staining (Fig2A). We hypothesized that the low hazy signal stains inactive domains, which have been previously shown to contain low levels of H3K27me3, while the high punctate signal labels the highly H3K27me3-enriched PcG domains (see confirmation below). Knockdown or mutation of PRC1 components *Pc* (Fig. S1A), *Ph*, or *Sce* (Fig. S1B) did not significantly affect H3K27me3, nor did mutation of PRC2 accessory subunit, *Jarid2* (Fig2A). However, *E(z)^GLKD^* completely removed the punctate and hazy signals, *Scm^GLKD^* removed most of the punctate signal while leaving the hazy signal intact, and *Pcl^GLKD^* lessened the intensity of both the punctate and hazy signals. Only *E(z)^GLKD^* affected H3K27me2 staining in stage 5 nurse cells (Fig2A). Thus, Scm likely promotes high H3K27me3 enrichment in PcG domains while not affecting the broad pattern of H3K27me2 or low H3K27me3 staining found on inactive domains. Although they do not block follicle progression after the onset of nurse cell differentiation, *Scm^GLKD^* females are sterile.

**Fig 2:**
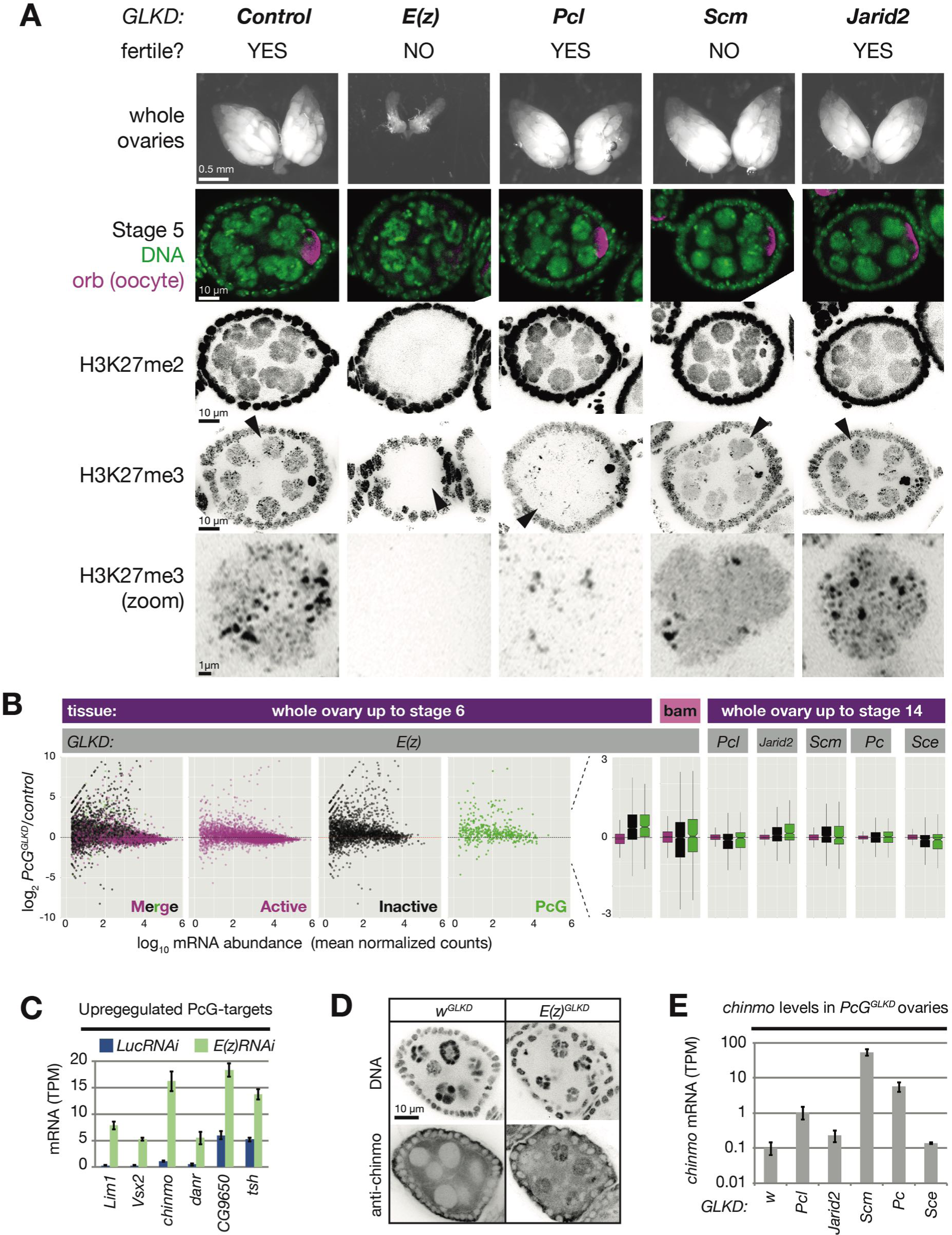
PRC2 represses endogenous genes in inactive and PcG domains in nurse cells (A) *PcG^GLKD^* effects on fertility, ovarian development, and bulk H3K27 methylation. Whole ovaries (row 1) or immunofluorescence (IF) images of stage 5 follicles antibody stained for the indicated protein epitopes or DNA (DAPI) (rows 2-5). *E(z)^GLKD^* blocks oocyte differentiation (row2), and abolishes H3K27me2 and H3K27me3 staining, while *Pcl^GLKD^* and *Scm^GLKD^* reduce H3K27me3 but not H3K27me2. (B) Whole ovary RNAseq showing how indicated *PcG^GLKDs^* affect median gene expression in whole ovaries that fully developed (right), developed until Stage 6 (left), or failed to differentiate due to the *bam* mutation (middle). Each dot represents a protein-coding gene colored to match the chromatin domain it resides in. Notched boxplots (right) summarize the fold-change distribution for all genes residing in each domain class. Notches show 95% confidence interval of the median, boxes show interquartile range. (C) Mean transcripts per million (TPM) of transcription factors located in PcG domains with the highest expression (> 5 TPM in *E(z)^GLKD^*) and upregulation (> 2.5-fold change) in *E(z)^GLKD^* compared to control *Luc^GLKD^*. Error bars represent 1 standard deviation from mean. (D) Stage 5 follicle IF showing Chinmo protein upregulation in *E(z)^GLKD^* nurse cell nuclei. DNA = DAPI. (E) Whole ovary RNAseq showing that GLKD of some *PcG* genes upregulate *chinmo*, compared to control (*w*). Size bars: A (row 1) 0.5 mm, (rows 2-4) 10μ, (row5) 1μ

Because Scm only affected H3K27me3 on PcG domains, we hypothesized that Scm enhances silencing on PcG domains while not affecting inactive domains. Consistent with this, *Scm^GLKD^* partially alleviated PcG but not inactive reporter silencing in nurse cells (Fig1I). For all three PcG reporters tested, *E(z)^GLKD^* more completely blocked silencing than *Scm^GLKD^*, suggesting that Scm enhances PRC2-dependent repression in PcG domains in nurse cells (Fig1H versus I). These results argue that PRC2 dependent repression acts on both inactive and PcG chromatin by a process that does not require H3K27me3, and complete repression in PcG domains requires Scm-dependent H3K27me3 enrichment.

### PRC2 suppresses endogenous gene expression within PCG and inactive domains

We next carried out experiments to investigate the effects of removing PcG proteins on endogenous gene expression in both nurse cells and their progenitors. Based on evidence from the reporter genes, we expected that *E(z)^GLKD^* would increase the expression of genes in both inactive and PcG domains in nurse cells but not progenitor cells (Fig. 2B). To investigate the consequences of *E(z)^GLKD^* in nurse cells, we extracted and sequenced mRNA from *control Luc^GLKD^* and *E(z)^GLKD^* whole ovaries containing equivalent stage distribution and used DEseq2 (Love et al., 2014) to quantify the effect of *E(z)^GLKD^* on protein-coding genes residing in different domain types. We additionally examined the effect of *E(z)^GLKD^* on progenitor cells by purifying mRNA from control or *E(z)^GLKD^* ovaries where germline stem cell differentiation is blocked by the *bag of marbles (bam)* mutation (Kai et al., 2005; McKearin and Spradling, 1990). In whole ovaries containing up to stage 6 nurse cells, *E(z)^GLKD^* had no effect on the median expression of genes in active domains while inducing a subtle, 1.4 fold median upregulation of genes in both inactive and PcG domains. In *bam* ovaries, containing GSC-like germ cells, *E(z)^GLKD^* had no effect on the median gene expression in any domain (Fig2B). Thus, PRC2 silences endogenous genes (as well as reporters) in both inactive and PcG domains in nurse cells but not progenitors.

Our observations that Pcl and Scm, but not most other PcG proteins, influence H3K27me3 levels suggested that loss of these genes might affect silencing at PcG domains but not inactive domains. We knocked down various PcG proteins in nurse cells (FigS2) and assayed their general effects on gene expression in different domain types (Fig. 2B). Surprisingly, we did not detect an increase in median gene expression in PcG domains following GLKD of *Pcl, Jarid2, Pc,* or *Sce*. Additionally, we did not detect a significant increase in median gene expression in *Scm^GLKD^*, which partially relieved reporter silencing in PcG domains. We hypothesized that PRC2 is sufficient to silence most genes in PcG domains through an H3K27me3/PRC1-independent mechanism, but that a subset of PcG domains require additional H3K27me3/PRC1-dependent silencing if strong activators specific to those domains are present.

We identified the strongest upregulated genes in PcG domains following *E(z)^GLKD^* in nurse cells in order to identify the potential biological targets of Polycomb repression. Six transcription factors were upregulated least 2.5-fold and expressed at more than 5 transcripts per million (TPM) in *E(z)^GLKD^* ovaries (Fig2C). The most highly upregulated such gene was *chinmo*, a Jak/Stat signaling target that is normally expressed in both male and female germline stem cells, plays an important role in male germ cell development, and can cause sex transformation in female cells (Flaherty et al., 2010; Grmai et al., 2018; Ma et al., 2014). We used anti-Chinmo antibodies to confirm that *E(z)^GLKD^* induced high Chinmo levels in nurse cell but not follicle cell nuclei (Fig. 2D). Finally, we tested whether other PcG proteins are required for *chinmo* silencing. We observed *chinmo* upregulation in *Pcl^GLKD^*, *Pc^GLKD^*, and *Scm^GLKD^* compared to control *w^GLKD^* (Fig2E). Thus, a variety of PcG proteins enhance silencing at this particularly vulnerable PcG domain in nurse cells. In conclusion, our experiments with endogenous and reporter genes show that canonical Polycomb silencing is absent from germline stem cells and initiates in nurse cells as they differentiate from progenitors and oocytes. It is likely that canonical Polycomb silencing acts during nurse cell differentiation to suppress a relatively small number of targets in nurse cells, including *chinmo*, that contain strong enhancers in the context of germline development.

### CBP increases and PRC2 changes targeting at the onset of PRC2-dependent silencing

We hypothesized that two general mechanisms could explain why E(z)-dependent silencing arises on both inactive and PcG domains as nurse cells differentiate. First, PRC2 activity, or other cofactors collaborating with PRC2 activity, might increase in differentiating nurse cells to induce silencing. Alternatively, regional PRC2 activity may remain constant, while an activator that opposes PRC2 activity increases. *nejire*, the fly homologue of *CBP/p300*, opposes PRC2 and activates transcription by catalyzing H3K27 acetylation (Holmqvist et al., 2012; Lee et al., 2015; Pasini et al., 2010; Tie et al., 2003) and promoting PolII recruitment or elongation through the +1 nucleosome of many genes including Hsp70 (Boija et al., 2017). *nejire* was previously isolated in a screen for ovarian defects (Yan et al., 2014), and we observed that *nejire^GLKD^* ovaries contained long chains of follicles that failed to grow significantly larger than stage 1 or 2 (Fig3A). Because *nejire^GLDK^* follicles are produced at a seemingly normal rate, but fail to grow after the onset of nurse cell differentiation, we hypothesized that nejire activity increases in differentiating nurse cells. To detect general changes in CBP activity, we stained ovaries for H3K27ac. *nejire^GLKD^* depleted H3K27ac antibody staining, confirming the specificity of the antibody for detecting nejire activity (Fig3A). As predicted, the intensity of H3K27ac staining in wild type ovaries increased dramatically following nurse cell differentiation (Fig3B). This increased nejire activity may be partly transcriptionally controlled, because we detected a doubling of *nejire* transcripts in differentiated cells versus GSCs (Fig3F). Thus, developmental upregulation of *nejire* may contribute to growing differences in reporter inducibility in active versus inactive domains as nurse cells differentiate. However, reporters in inactive domains were more strongly induced in GSCs and region 1 cysts, which have low *nejire* activity, than in late stage nurse cells, which have high *nejire* activity (Fig1 G), indicating that PRC2-dependent silencing also increases in nurse cells.

The increased PRC2-dependent silencing in differentiating nurse cells might result from increased total PRC2 activity or from changes in PRC2 targeting. Unlike *nejire*, *E(z)* and other core PRC2 subunits were present at similar mRNA levels in GSCs and differentiated cells (Fig 3F, Fig 5B), and the levels of H3K27me2, the most abundant product of PRC2 activity, did not significantly change during germline development (Fig 3D). However, H3K27me1 levels increased, and H3K27me3 distribution dramatically changed as PRC2-dependent silencing appeared in nurse cells (Fig. 3C,E). In GSCs and pre-meiotic germ cells in region 1 of the germarium, H3K27me3 staining appeared diffuse compared to the condensed H3K27me3 foci found in somatic follicle cells surrounding the germline (Fig3E,H). In region 2, as germ cells enter a prolonged pre-meiotic S-phase and prophase arrest, H3K27me3 levels increased and small areas of signal enrichment began to form (Fig. 3E). As follicles leave the germarium and nurse cells begin to endocycle, H3K27me3 puncta became more pronounced in nurse cells while non-punctate chromatin staining decreased. However, oocytes, which are arrested in meiotic prophase as nurse cells grow, retained a high and diffuse H3K27me3 signal (Fig3E). We additionally noted a similar dichotomy in H3K27me3 staining between pre-meiotic germ cells and surrounding somatic follicle (granulosa) cells in the E14.5 fetal mouse ovary – a time when germ cells are at similar developmental stages as germ cells in region 2a of the fly germarium (Fig3G,H). These data imply that the transition from a diffuse H3K27me3 staining pattern in pre-meiotic germ cells and oocytes to a punctate pattern associated with increased silencing in differentiating nurse cells and somatic cells reflects a developmentally programmed change in PRC2 distribution rather than overall increased PRC2 activity.

**Fig 3:**
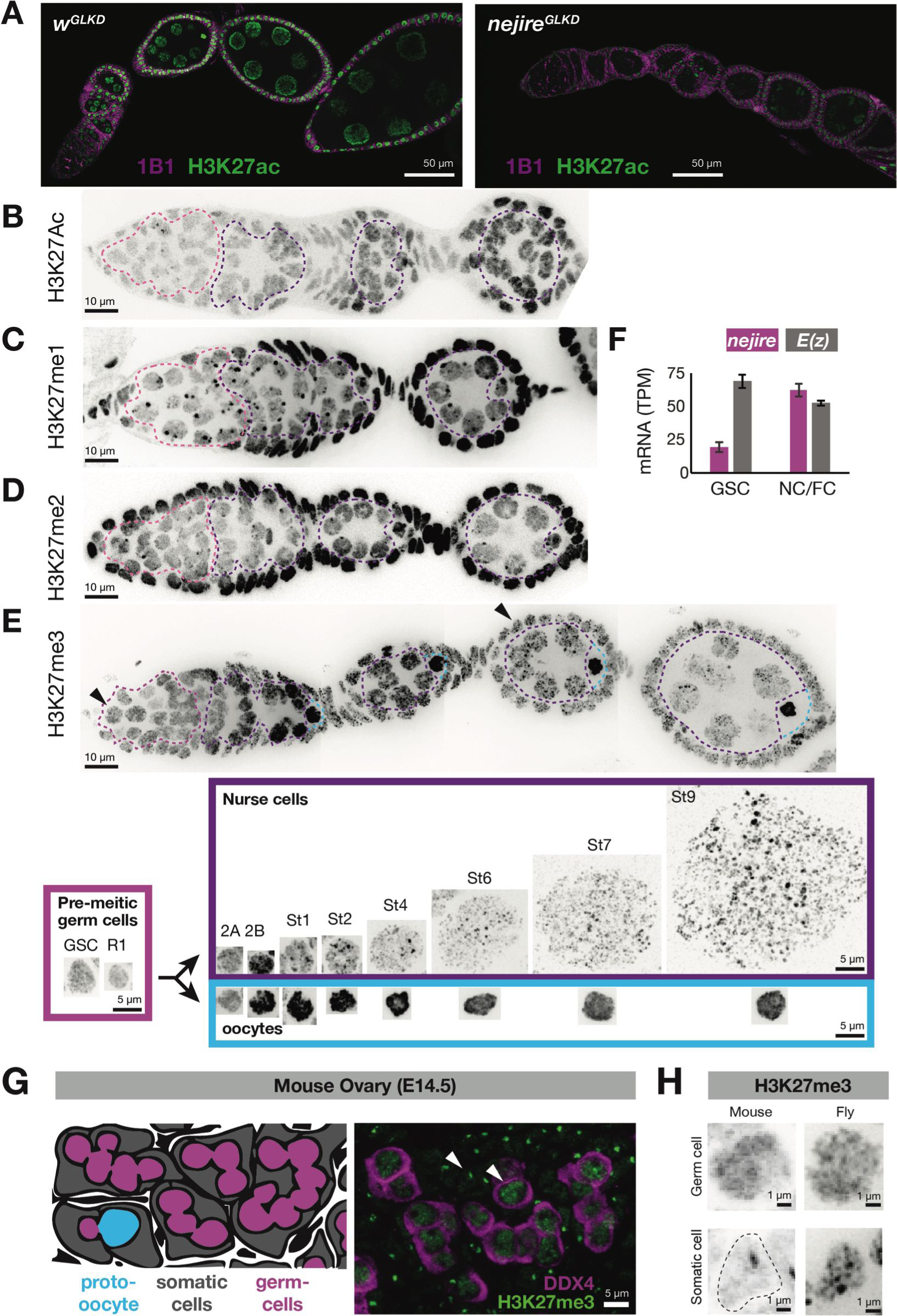
H3K27acetylation and methylation changes during nurse cell differentiation (A) IF staining showing *nejire* germline knockdown (*nej^GLKD^*) reduces H3K27ac in germ cells and arrests follicle development. (B-D) IF staining of H2K27Ac (B), H3K27me1 (C), H3K27me2 (D), or H3K27me3 (E). Summary below shows H3K27me3-stained nuclei (stages indicated) from premeiotic germ cells (pink box), nurse cells (purple box) or oocytes (cyan box). (F) *nejire* and *E(z)* mRNA levels (TPM) in FACS purified GSCs or whole ovaries comprised mostly of differentiated nurse and follicle cells (NC/FC). (G) Cell diagram depicting an E14.5 mouse ovary (left) showing germ cell cysts (magenta), a proto-oocyte (cyan), all surrounded by somatic cells (grey). Right: E14.5 mouse ovary (IF) stained for DDX4 (germ cells: magenta) and H3K27me3 (green). (H) H3K27me3 staining of *Drosophila* and mouse nuclei (arrowheads in E and G, respectively) showing similar diffuse signal in progenitor germ cells and punctate signals in differentiated follicle cells. Scale bars: A 50μ, B–E 10μ, E lower 5μ, F 10μ. G 10μ.

### Mapping differences in PRC2 localization before and after nurse cell differentiation

To understand how the H3K27me3 pattern changes during nurse cell differentiation, we mapped the positions of H3K27me3 nucleosomes throughout the genomes of GSCs and nurse cells using chromatin immuno-precipitation and deep sequencing (ChIPseq) (Fig4). We used nuclear fractionation and a fluorescence activated cell sorter to purify fixed, GFP or Tomato-labeled nuclei from a variety of cell types in the ovary. We purified G1 germline stem cells from *bam* mutant ovaries, where they are highly enriched, using a germline driven NLS-GFP and G1 (2c) DAPI content as tags (Fig4A). For nurse cells and follicle cells, we used germline driven tdTomato and DAPI to purify individual nurse cell and follicle stages beginning at 2c and ending at 512c DNA content (Fig4B). For all of our ChIP experiments, we used a constant amount of input chromatin and either a mouse or alternative fly species spike-in to normalize relative amounts of H3K27me3 between samples.

ChIPseq comparison of germline stem cells to differentiated nurse cells or follicle cells revealed a dramatic change in the distribution of H3K27me3, consistent with the immunofluorescence staining. Relative to spike-in, germline stem cells had about 30% more H3K27me3 signal than nurse cells or follicle cells. However, GSC H3K27me3 was evenly distributed in a non-canonical pattern while nurse cell and somatic follicle cell H3K27me3 was highly enriched on canonical PcG-target loci (Fig4C). We quantified the non-canonical H3K27me3 enrichment pattern by dividing the genome into overlapping 5kb bins and classifying the bins according to a modEncode chromatin state model from S2 cells. In germline stem cells, we noted no greater enrichment of H3K27me3 on PcG domains compared to generally inactive domains, in contrast to an approximately 10-fold enrichment in differentiated cells (Fig4E,F). H3K27me3 signal in GSCs was significantly depleted from modEncode-mapped active domains (Fig4F) and from the gene bodies of genes with H3K27ac transcription start sites (TSSs) (Fig4D). While GSCs and nurse cells had similarly low H3K27me3 enrichment on active domains, nurse cells had significantly less H3K27me3 on inactive domains and more H3K27me3 on PcG domains than GSCs (Fig.4F). Thus, GSCs have a non-canonical distribution of H3K27me3 that appears to be redistributed from inactive to PcG domains during nurse cell differentiation to produce in a canonical H3K27me3 pattern common to somatic cells.

**Figure 4:**
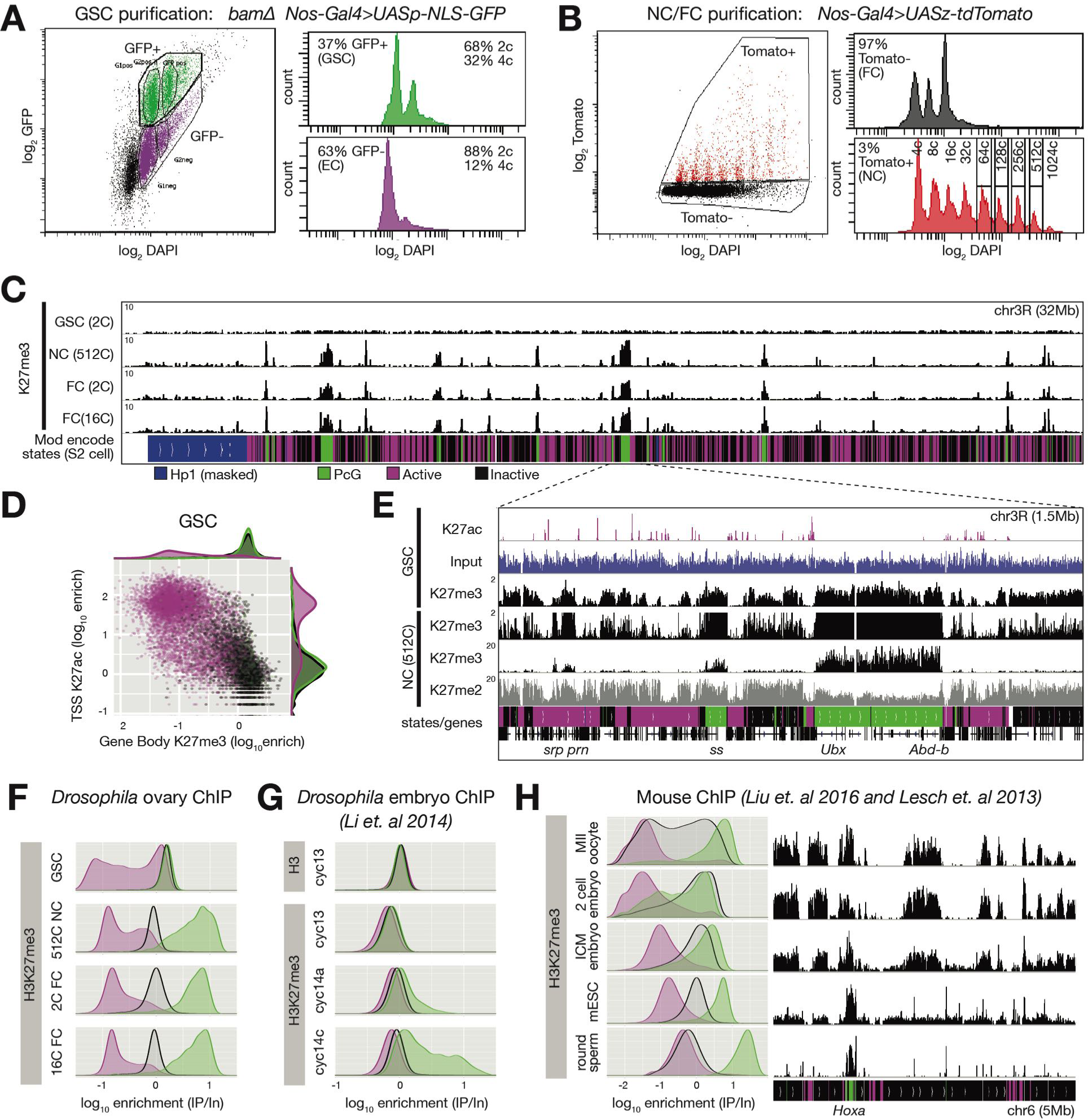
Germline stem cells and early embryos have non-canonical H3K27me3 (A,B) FACSdiva-generated summaries of FACS-sorted fixed nuclei. (A) Nuclei sorted in two steps using GFP from *bam* ovaries expressing germline-specific nuclear GFP vs DNA content (DAPI), yielding GSCs and somatic escort cells (EC). (B) Nurse and Follicle cell nuclei sorted in two steps using Tomato from ovaries expressing germline-specific tdTomato vs. DNA content (DAPI). The haploid DNA content (C-value) is noted above each peak. 2c nuclei were not on scale to aid visualization of larger nurse cells. (C) Chromosome 3R genome browser view of spike-in normalized H3K27me3 ChIPseq read depth from the indicated purified nuclei. Below: chromatin states in S2 cells. (D) H3K27ac enrichment (IP/Input) in a 500bp bin downstream from annotated transcription start sites (TSS) vs. corresponding H3K27me3 gene body enrichment (IP/Input). (E) Chromosome 3R subregion (dashed lines) showing spike-in normalized read depth from input (blue) or ChIPseq of the indicated epitopes. Nurse cell H3K27me3 read depth is plotted on two scales to show enrichment over a 100-fold range. (F-H) H3K27me3 enrichment histograms across active (magenta), inactive (black), and PcG (green) domains in 5kb bins for the indicated fly and mouse cell types (see Methods). (G) H3K27me3 ChIP data from Drosophila cycle 13 and 14 embryos (Li et al., 2014) plotted in the same manner. (H) H3K27me3 ChIP data showing a region around HoxA from preimplantation mouse embryos and embryonic stem cell cultures (Liu et al., 2016) or differentiated round spermatids (Lesch et al., 2013) (right), and summarized in the same manner as (F,G) (left).

The non-canonical H3K27me3 distribution we observed in germline stem cells resembled the non-canonical H3K27me3 distribution previously reported in mouse MII arrested oocytes and preimplantation embryos (Liu et al., 2016; Zheng et al., 2016). To compare the non-canonical H3K27me3 distributions between species, we categorized the mouse genome into active, inactive, and PcG domains (see Methods) and reanalyzed mouse oocyte and preimplantation embryo H3K27me3 ChIPseq (Fig4H). In oocytes and preimplantation mouse embryos, like in fly GSCs, H3K27me3 was depleted from active loci and present at similar levels on inactive and PcG domains. Thus, both Drosophila germ cells and mouse embryonic somatic cells transition from a common non-canonical H3K27me3 distribution in precursors to a canonical H3K27me3 distribution during differentiation.

We studied early fly embryos to see if they undergo similar changes in H3K27me3 distribution as part of their well-characterized differentiation program. During the rapid initial 13 embryonic divisions in fly embryos, most histones are recently deposited in S-phase and contain low levels of post-translational modifications. However, at cycle 14, the cell cycle dramatically slows and many histone modifications become apparent by antibody staining. By ChIP, H3K27me3 enrichment on PcG domains begins at cycle 14, with much weaker signal in cycle 13 chromatin (Li et al., 2014). To determine whether total PRC2 activity or targeting changes from cycle 13 to cycle 14, we re-analyzed the *Li et al.* libraries with our genome segmentation strategy. At cycle 13, H3K27me3 was distributed in a non-canonical pattern resembling GSCs and mouse embryos, with equal enrichment on inactive and PcG domains and a slight depletion on active domains (Fig4F). Thus, fly embryos also change from a non-canonical to a canonical H3K27me3 distribution at the onset of somatic differentiation in cycle 14.

### Core PRC2 subunits remain constant while PRC1 shows little activity in nurse cells

To investigate how the transition from a non-canonical to a canonical H3K27me3 distribution is brought about, we first looked for changes in the levels of PcG RNAs and proteins that might affect the specificity of PRC2 (Fig. 5). Consistent with stable core PRC2 mRNA levels (Fig. 3F, 5B), fluorescence microscopy showed that the PRC2 core component, Su(z)12, did not change significantly in abundance even as it coalesced into perinuclear puncta in differentiated nurse cells (Fig. 5D). Jarid2 was strongly elevated in somatic follicle cells, consistent with the observed increase via RNAseq, but it showed only a small increase during nurse cell differentiation (Fig. 5B,D). Utx and Asx/Calyposo mRNA, which encode enzymes opposing PRC2 and PRC1 activities, respectively, decreased 2-3 fold (Fig. 5B). mRNA encoding the PRC1 components Psc and Sce decreased 2-4 fold, while Pc mRNA and protein, and Ph protein, increased slightly and formed perinuclear puncta in nurse cells (Fig. 5B,D). The enzymatic product of PRC1 activity, H2AK119ub was nearly undetectable in germ cells until stage 7, unless we depleted Asx, a subunit of the, H2AK119 deubiquitylase (FigS3). In *Asx^GLKD^*, H2AK119ub was diffusely localized in GSCs and formed prominent puncta after nurse cell differentiation, similar to PRC1 components. In summary, PRC1 activity remained at a low level, while two PRC1 subunits, Pc and Ph, slightly increased in abundance and localized to PcG domains as nurse cells differentiated.

**Fig5.**
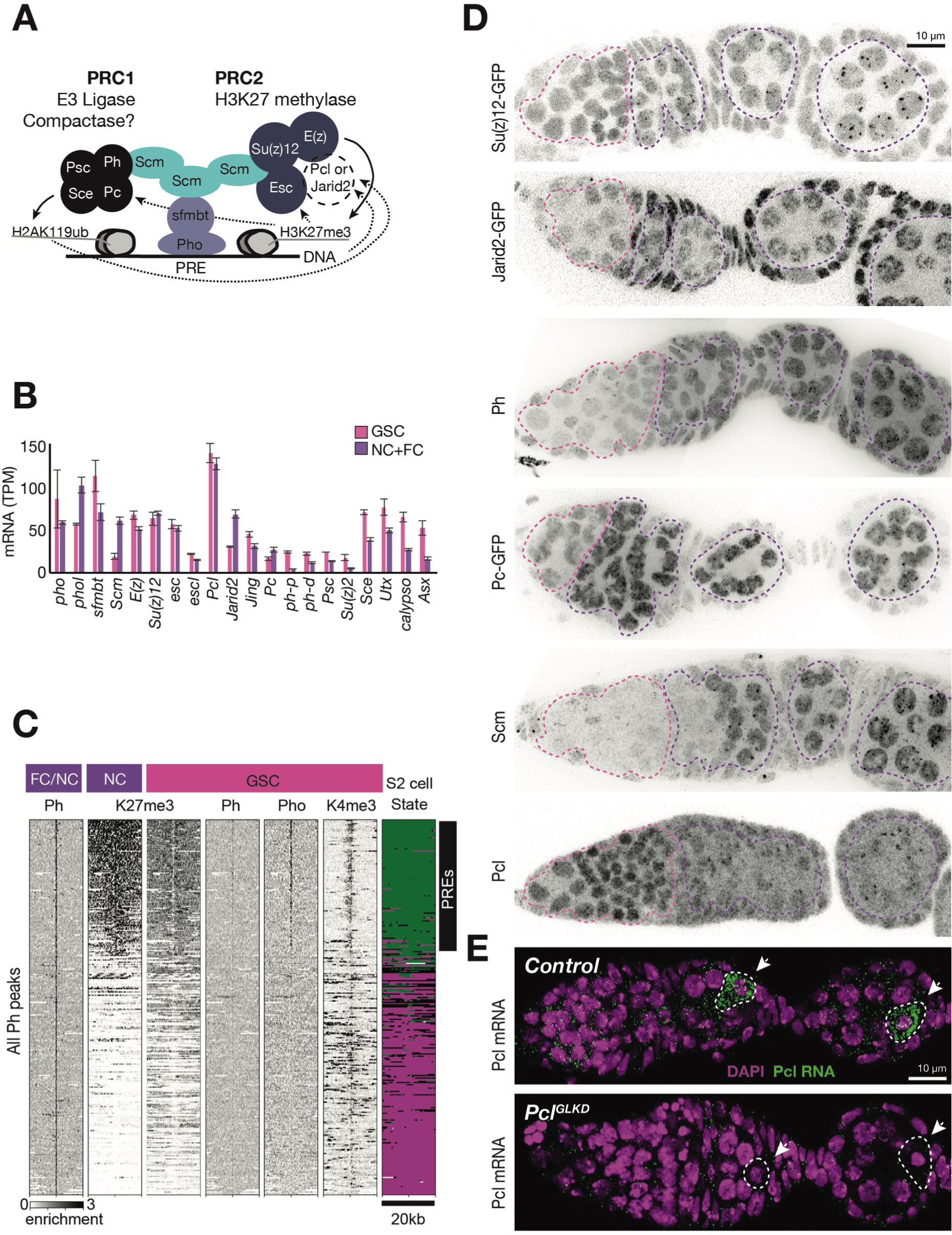
PcG-gene expression and localization during germline development (A) Model showing subunits and activities of PRC1, PRC2, PhoRC, and Scm, a putative bridge between complexes. (B) PcG gene mRNA levels (TPM) measured by RNAseq analysis of FACS-purified GSC (pink) vs whole ovary tissue enriched in differentiated nurse and follicle cells (purple, NC+FC). (C) ChIPseq raw read depth heatmap comparing PcG proteins or histone modifications in 20kb regions surrounding every Ph peak found in differentiated ovary tissue. In GSCs, Ph peaks in PcG domains (upper region) are associated with Pho and a “bivalent” enrichment of H3K27me3 and H3K4me3. (D) IF (for Ph, Scm, and Pcl) or native GFP fluorescence (of GFP-tagged Su(z)12, Jarid2, or Pc) showing developmental changes of the indicated PcG proteins from region 1 (pink lines) containing progenitors, through region 2a and early follicles (purple lines). (E) *In situ* hybridization shows that *Pcl* mRNA (green) accumulates in oocytes (arrowheads, white outline). *Pcl^GLKD^* serves as a control. Scale bars: D,E 10μ.

### GSC chromatin resembles mammalian progenitor chromatin and exhibits bivalent domains

Because GSCs expressed, but did not localize Ph into perinuclear puncta, we used ChIP-seq to determine whether Ph or its recruitment factor, Pho (Brown et al., 2018) were bound to PREs in GSCs (Fig 5C). GSCs lack Polycomb silencing, suggesting that the PREs within them are inactive. Surprisingly, GSC PREs bound Pho (Fig 5C), and Pho-bound PREs were moderately enriched for both H3K27me3 and H3K4me3- the “bivalent” chromatin signature frequently observed in mammalian progenitor cells. However, despite the presence of Pho, we detected far less Ph binding to PREs in GSCs than in differentiated cells (Fig.5C). We conclude that PREs are engaged by Pho in GSCs but are unable to recruit other PcG proteins.

### Pcl and Scm regulate PcG domain formation in developing nurse cells and oocytes

We identified two developmentally regulated PcG proteins that could initiate the targeting of PRC1 and PRC2 to Pho-bound PREs upon nurse cell differentiation. Scm, which potentially directly links PRC1 and PRC2 to PhoRC (Fig. 5A), was dramatically upregulated in region 2B nurse cells (Fig. 5D). Scm induction correlated with the formation of nuclear foci of H3K27me3 (Fig. 3E), and of PRC1 and PRC2 subunits (Fig5D). Without Scm, the majority of these foci never formed in the germline (FigS1A,C).

Pcl represents the second PcG protein whose expression changes during nurse cell differentiation. Pcl is present at high levels in GSCs and it is required for the formation of intense H3K27me3 puncta in nurse cells (Fig2A). However, Pcl levels sharply decrease on average in early stages of nurse cell differentiation (Fig.5D). We performed fluorescence *in situ* hybridization for Pcl mRNA and found that Pcl mRNA dramatically changes its localization at this time (Fig. 5E). Rather than remaining in all cells, Pcl mRNA accumulates in the oocytes of follicles by stage 1, while falling substantially in nurse cells (Fig.5E). We also investigated whether Pcl protein levels similarly decrease during somatic cell differentiation in cycle 14 embryos. We noted an abrupt decrease in maternally expressed Pcl-GFP fluorescence at cycle 14, suggesting that Pcl gene products are degraded at the maternal to zygotic transition (Fig.S4). Therefore, in both the ovary and embryo, Pcl levels inversely correlate with the formation of canonical PcG domains in differentiating cells.

### Analysis of mutant chromatin suggests a model for how Pcl and Scm initiate Polycomb silencing

To better understand how changes in Pcl and Scm modulate Polycomb silencing during nurse cell differentiation, we investigated how these genes influence H3K27me3 levels by ChIP-seq (Fig. 6A-E). To highlight changes at PRE sites, we subdivided PcG domains into 5kb bins either containing (Fig6B, orange), or lacking (Fig6B, green) a PRE. Scm is not significantly expressed in GSCs, but depleting Scm in nurse cells reduced H3K27me3 enrichment on PRE and PcG domains but not on inactive domains. Thus, Scm depletion resulted in a nearly equal enrichment of H3K27me3 on inactive, PcG, and PRE domain types (Fig. 6A,D). Thus, without Scm function in differentiating nurse cells, H3K27me3 remained in a non-canonical pattern similar to wild type GSCs or early fly and mouse embryos.

**Figure 6:**
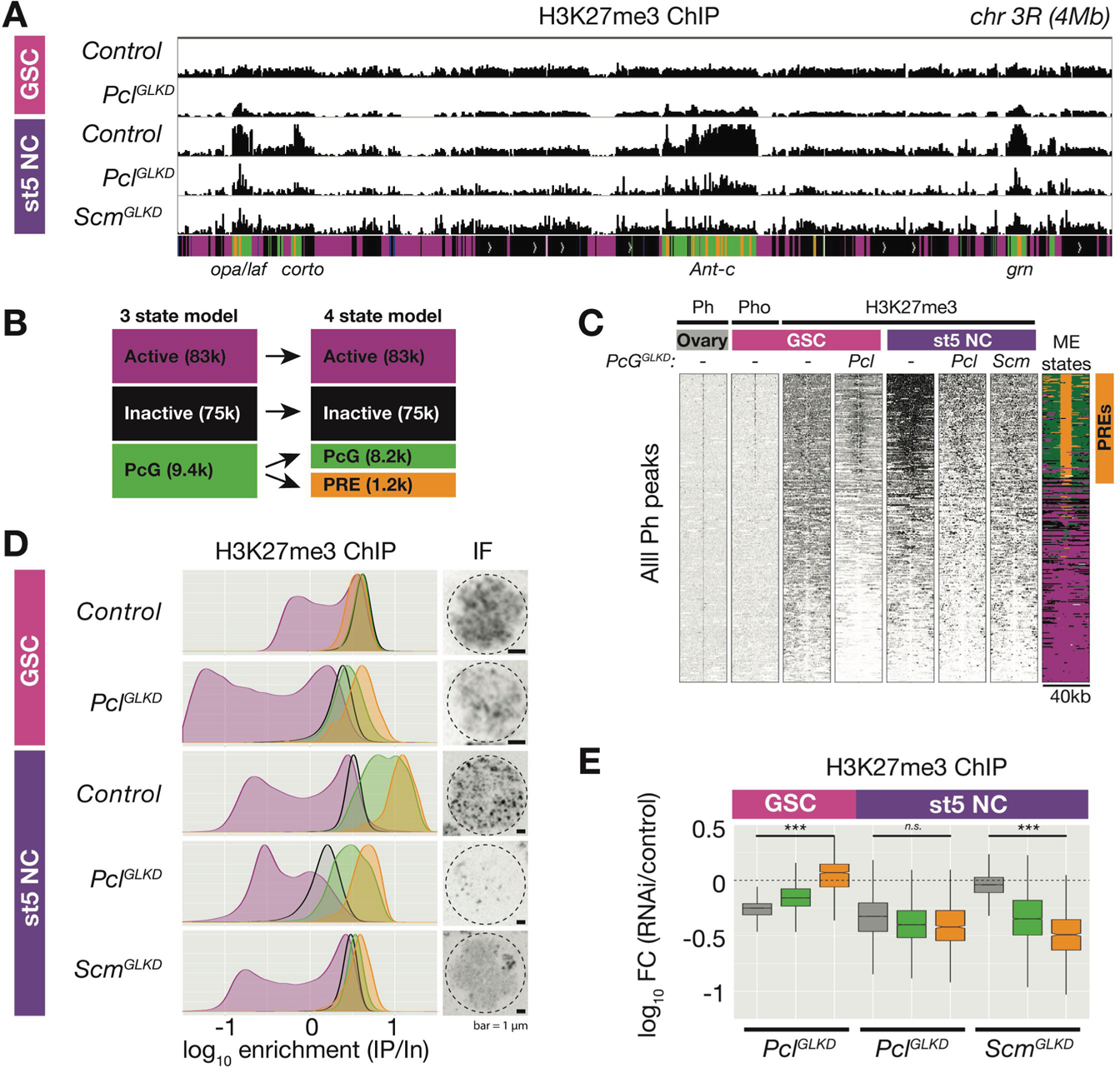
The effects of Pcl and Scm on H3K27me3 domain formation (A) Spike-in normalized H3K27me3 ChIP from FACS-purified GSC or stage 5 nurse cell (St5 NC) nuclei of the indicated genotypes in a 4Mb region including *Ant-c*. *Pcl^GLKD^* in GSCs specifically depletes H3K27me3 from inactive loci. In NCs, *Pcl^GLKD^* and *Scm^GLKD^* deplete H3K27me3 from PcG loci. (B) Subdivision of 9,400 PcG bins into 1,200 PRE-containing bins (orange), and 8,200 PRE-lacking bins (green). (C) ChIPseq raw read depth heatmap showing the effect of *Pcl* and *Scm* knockdown on H3K27me3 enrichment near all ovary Ph peaks. Note that H3K27me3 enrichment on GSC PREs is revealed by *Pcl^GLKD^.* (D) Smoothed histograms for each indicated genotype and stage showing spike-in normalized H3K27me3 enrichment (IP/Input) in 5kb active (magenta), inactive (black), PcG (green), and PRE-containing (orange) bins tiling the genome. IF images of H3K27me3 in GSC and St5 NC show the relationship between H3K27me3 enrichment in ChIPseq (left) and whole mount staining (right). (E) Boxplots summarizing the fold changes in H3K27me3 enrichment in inactive (black), PcG (green), and PRE-containing (orange) 5kb bins induced by *Pcl^GLKD^* or *Scm^GLKD^* in the indicated stages.

In nurse cells, *Pcl^GLKD^* reduced H3K27me3 enrichment equivalently from inactive, PcG and PRE-containing bins (Fig6D). In contrast, in GSCs, *Pcl^GLKD^* disrupted the normally even distribution of H3K27me3 across PcG and inactive domains, by upregulating H3K27me3 enrichment on PcG domains and PREs relative to inactive domains (Fig.6A,C,D). Genome-wide, PRE-containing bins showed a median 1.2 fold increase in H3K27me3 enrichment following Pcl knockdown, while inactive domain bins showed a 1.8 fold decrease in H3K27me3 enrichment (Fig.6E). This strongly suggests that high Pcl levels promote the non-canonical H3K27me3 characteristic of progenitors.

Because *Pcl^GLKD^* mostly affected inactive domain H3K27me3 in progenitors (Fig 6E), we wondered whether *Pcl^GLKD^* also affects the abundance of H3K27me1 and me2– the most common H3K27 methylation states in inactive chromatin in somatic cells (Kharchenko et al., 2011). We performed antibody staining on *control*, *Pcl^GLKD^* and *E(z)^GLKD^* ovaries and measured antibody staining intensity in the euchromatic regions of individual GSCs (Fig. S6). All three antibodies labeled E(z)-dependent methylation because *E(z)^GLKD^* completely eliminated chromatin staining in germ cells. Consistent with ChIPseq, *Pcl^GLKD^* depleted H3K27me3 staining by 2.3-fold (Fig.7A). However, *Pcl^GLKD^* increased H3K27me2 and H3K27me1 abundance by 1.4-fold and 14-fold, respectively (Fig7A). Therefore, in GSCs, high Pcl levels promote H3K27me3 while reducing the abundances of lower H3K27 methylation states.

**Figure 7:**
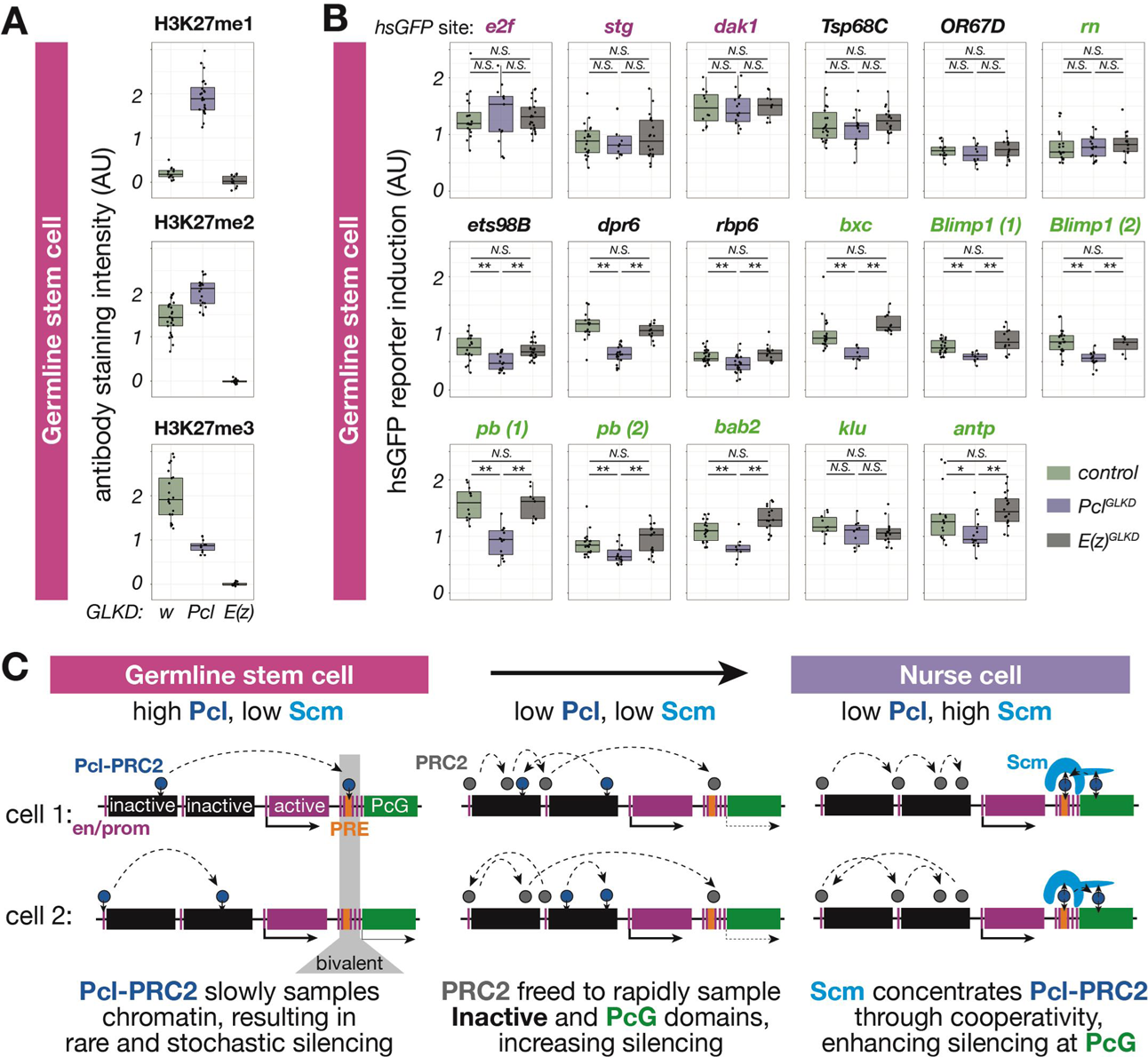
Pcl controls PRC2 sampling rate and silencing in GSCs (A) Quantification of relative H3K27me1/2/3 antibody staining intensity in the euchromatin of control *w^GLKD^*, *Pcl^GLKD^*, or *E(z)^GLKD^* GSCs. (B) Quantification of reporter gene induction in GSCs in control *w^GLKD^*, *Pcl^GLKD^*, or *E(z)^GLKD^*. Note that *Pcl^GLKD^* reduces the induction of some inactive (black) and PcG (green) localized reporters but not active (magenta) localized reporters (* = p<0.05, ** = p<0.01, N.S.=not significant; Student’s t-test, unpaired, 2-tailed). (C) PRC2 sampling model for the developmental control of silencing. In GSCs, most PRC2 is associated with Pcl (dark blue), and samples a small number of different sites in different cells, resulting in infrequent and stochastic silencing of regulatory regions (magenta and orange). As Pcl levels drop during differentiation, core PRC2 (grey) is freed to sample and silence more sites. As Scm (light blue) is induced and concentrated on PREs (orange), it preferentially concentrates Pcl-PRC2 through cooperativity, and locally increases PRC2 sampling and silencing.

Because of its dramatic affect on H3K27 methylation state, we tested whether *Pcl^GLKD^* affects Polycomb silencing in GSCs. We measured the induction of 5 reporters in inactive domains, 9 in PcG domains, and 3 in active domains, and found that *Pcl^GLKD^* significantly suppressed reporter induction in 3/5 inactive, 7/9 PcG domains, and 0/3 active domains (Fig7B) and did not enhance induction in any domain. Thus in GSCs, Pcl inhibits, rather than promotes Polycomb silencing. This result was particularly surprising because Pcl promotes H3K27me3 in these cells, and H3K27me3 is generally thought to enhance silencing. To explain this paradox, below we propose a model for Polycomb silencing that invokes PRC2 sampling rate rather than the methylation state of PRC2 modified domains.

## Discussion

### The Drosophila germline as a model for chromatin silencing

Nurse cells are found in diverse animal species, but they have traditionally been considered germ cells rather than late differentiating somatic cells. Here we show that after separating from the oocyte lineage, *Drosophila* nurse cells modulate PcG protein expression, and enrich H3K27me3 locally to silence PcG domains, much like embryonic somatic cells.

Moreover, germline stem cells, the precursors of nurse cells and oocytes, lack gene silencing and contain a non-canonical distribution of H3K27me3, like pluripotent cells in mouse and *Drosophila* embryos (Liu et al., 2016; Zheng et al., 2016)(Fig. 4H). These findings are fundamentally important for understanding oocyte and nurse cell biology and indicate that nurse cells undergo a differentiation process indistinguishable from embryonic somatic cells.

The Drosophila female germline offers many advantages for studying the biology of progenitor cell chromatin, and for understanding chromatin silencing during cell differentiation. Female GSCs continuously divide to produce new progenitors and nurse cells throughout life, generating a vastly greater mass of cellular material than all other adult tissues combined. GSCs are easily expanded by mutation, and nurse cells become progressively more polyploid, allowing large quantities of chromatin to be purified for molecular genomic analyses from each developmental stage. Custom transgenic reporters can be targeted to diverse chromatin types throughout the genome and used to functionally test local chromatin properties. In the future, these combined genetic and biochemical methods have the potential to further advance our understanding of chromatin regulation during development.

### PRC2-mediated repression extends beyond PcG domains by regulated PRC2 targeting

The studies of female germ cells reported here revealed a clearer picture of Polycomb mediated repression and how it arises. GSCs, the progenitors of oocytes and nurse cells, exhibit a broad, non-canonical H3K27me3 distribution, along with substantial H3K27me2 modification of “inactive” chromatin domains that will eventually be repressed by PRC2, but reporters in both types of chromatin remain active. In contrast, differentiated nurse cells deplete H3K27me3 from inactive domains, enrich H3K27me3 on PcG domains, and silence reporters in both domain types. Thus, the initiation of silencing involves a major change in the distribution of PRC2 activity reflected by changes in H3K27 methylation, but is not generally associated with a particular H3K27 methylation state. Because our reporter is not regulated by transcription factors in traditional PcG domains, and our approach measures position-dependent silencing, we propose that PRC2 directly silences H3K27me3-depleted inactive domains in addition to PcG domains. Our work further supports the idea that PRC2 silences inactive domains through a random roaming mechanism (Ferrari et al., 2014; Lee et al., 2015; Pirrotta, 2016), and extends this model by showing that PRC2 roaming is developmentally regulated.

Several proteins or protein complexes have been proposed to influence PRC2 targeting through direct interactions, including Jarid2/AEBP2, Pcl, and Scm (Cooper et al., 2016; Kang et al., 2015; Li et al., 2010; 2017; Perino et al., 2018). In developing nurse cells, the change in PRC2 distribution on chromatin is associated with a substantial decrease in Pcl levels from their high abundance in GSCs, as well as a major increase in Scm expression.

### A PRC2 sampling rate model for Polycomb silencing establishment

By combining our results with previous biochemical studies of PcG proteins, we propose a sampling rate model for how PRC2-dependent silencing initiates during development. PRC2-dependent silencing at a particular regulatory site would depend on the relative frequencies with which PRC2 or transcriptional activators visit that site and not simply on its H3K27 methylation state. We propose that PRC2-interacting proteins regulate PRC2 sampling rate during development by modifying both the local concentration of PRC2 and its dissociation kinetics from chromatin. Pcl, a chromatin binding subunit of PRC2, triples PRC2’s residence time on DNA *in vitro* (Choi et al., 2017), and additional interactions between PRC2 and RNA, histones, and other chomatin-bound proteins including Scm could further lengthen the residence time of Pcl-PRC2 at a one site and impair its sampling rate and silencing ability at other sites. In principle, such changes in PRC2 persistence can explain the changes in PRC2 action that underlie Polycomb repression during differentiation.

In germline stem cells (Fig. 7B), abundant Pcl would saturate PRC2 and antagonize silencing by slowing PRC2 sampling. In support of this model, GSCs contained low levels of H3K27me1 – a product of fast sampling PRC2, and Pcl depletion dramatically increased H3K27me1 and reporter silencing and decreased H3K27me3 (Fig 7A,B, FigS7). Furthermore, by slowing PRC2 sampling and/or blocking its interaction with Jarid2, abundant Pcl would prevent H3K27me3 from accumulating around PREs and spreading throughout PcG domains. Instead, in each GSC, a random subset of PREs would be partially modified, and spreading would be inefficient and incomplete.

As Pcl levels drop and Scm levels increase during differentiation (Fig. 7C), two pools of PRC2, core PRC2 and Pcl-PRC2, would differentially promote silencing at inactive and PcG domains. At inactive domains, core PRC2 (freed of Pcl) would rapidly sample and silence regulatory regions through a random “roaming” mechanism. PRC2 might preferentially roam inactive domains or else PRC2 action on active domains might be overridden by compartmentalization of transcription-promoting factors (Corrales et al., 2017). At PcG domains, Scm would concentrate remaining Pcl-PRC2 through cooperativity due to Pcl-PRC2s interactions with both Scm and DNA. Thus, upon nurse cell differentiation, PRC2 sampling increases at both PcG domains (due to higher PRC2 concentrations) and at inactive domains (due to higher PRC2 mobility).

Why might PRC2 sampling be increased through two different mechanisms? Two mechanisms could allow inactive and PcG domains to be independently regulated. For example, a partly differentiated cell could maintain silencing in a Hox cluster while reducing silencing in inactive domains. Additionally, the enrichment-based mechanism at PcG domains could enhance silencing at particularly vulnerable sites by generating high levels of H3K27me3 and recruiting PRC1. This mechanism could limit the output of signal-induced transcription factors or silence enhancers with a high density of transcription factor binding sites such as the Hox clusters. For example, nurse cells, which are exposed to the JAK/STAT activator, Unpaired, highly over-expressed the JAK/STAT target *chinmo,* and the Hox cluster gene, *Abd-b* in *Scm^GLKD^* (Fig2E, FigS2). However, most genes in PcG domains were minimally affected by Scm and Pcl depletion compared to E(z) depletion, suggesting that roaming PRC2 is sufficient to silence most genes in nurse cells and additional Scm/Pcl-dependent silencing at PcG domains is differentially required in different cell types.

### The nature of the progenitor state

Stochastic gene expression in progenitors precedes numerous cell fate decisions in development (Urban and Johnston, 2018), and both PcG proteins and H3K27ac can contribute to stochasticity (Kar et al., 2017; Nicolas et al., 2018). Our work suggests that Pcl could generally regulate stochastic gene expression during development by regulating PRC2 sampling rate.

Slow sampling Pcl-PRC2 in progenitors might have a much lower probability of methylating a particular regulatory site compared to faster sampling core PRC2 in differentiated cells. Thus, in a population of progenitors, one would expect a mix of sampled (methylated) and not sampled (unmethylated) regulatory sites, which when averaged could generate the “bivalent” chromatin signature observed in progenitor chromatin.

Why would it be advantageous for GSCs and other progenitors to enhance gene stochasticity by slowing PRC2 sampling? By slowing, but not eliminating PRC2, GSCs could promote the developmental induction of previously inactive genes for differentiation but still be able to reverse the rare induction of inappropriate genes. Disrupting non-canonical PRC2 targeting may have subtle effects, for example *Pcl^GLKD^* might delay the transition from progenitor to differentiated cell while *E(z)^GLKD^* might enhance the appearance of rare, inappropriately differentiated cells. Although these effects would be difficult to detect, they may be enhanced by ectopically inducing *nejire* activity, which is normally low in GSCs.

### Implications for mammalian Polycomb silencing

The findings reported here suggest that the apparent differences in Polycomb silencing between Drosophila and mammals results primarily from differences in genome size. Animals with smaller genomes such as Drosophila may be able to effectively silence most enhancers in differentiated tissues though randomly roaming PRC2. However, small genomes may also contain regions sufficiently enriched in regulatory enhancers so as to require focused inhibition to ensure complete silencing. Drosophila PREs are embedded within large regulatory regions enriched for developmentally-regulated transcription factor binding sites (modENCODE Consortium et al., 2010). The larger number of PcG domains in Drosophila, compared to mammals may reflect the greater number of concentrated regulatory elements that require concentrated PRC2 activity. Nonetheless, the Drosophila genome contains a lot more inactive chromatin regulated by roaming PRC2 than PcG domain chromatin.

Roaming would likely be ineffective in the human or other large genomes, since PRC2 would non-productively roam large deserts of non-regulatory DNA. In mammals, PcG proteins silence hundreds of developmentally induced genes and PRC2 catalyzes high levels of H3K27me3 on CGIs clustered near their individual regulatory regions (Boyer et al., 2006; Bracken et al., 2006; Lee et al., 2006; Squazzo et al., 2006). The fly homologues of most mouse PRC2-targeted genes reside in inactive, rather than PcG domains (Fig S5). In mammals, enhancers are more widely distributed and only a few large clusters of regulatory elements, such as Hox loci, may require the inactivation of a large domain.

How, does PRC2 recognize a large number of diverse gene targets without the aid of PRE sequences? Concomitant with genome expansion, PRC2 might have evolved a preference for sites commonly appearing near regulatory regions. By preferring CG-rich sites in an organism utilizing CG methylation, the elevated mutation rate of these sequences might speed their removal from non-regulatory regions and focus PRC2 regulation. Transitions between random roaming and specific targeting have likely occurred many times during animal evolution. This may underlie the association between CGIs, CG methylation and large genome size, and the general inverse correlation between genome size and CG abundance (Lechner et al., 2013).

In mammals, Pcl proteins are one of several mechanisms proposed to target PRC2 to regulatory sites (Laugesen et al., 2019; Li et al., 2017; Perino et al., 2018). Removal of Pcl proteins from mESCs reduces H3K27me3 enrichment on CGIs without affecting low H3K27me3 levels outside of PcG domains, suggesting that the majority of Pcl-PRC2 complexes in mESCs specifically sample CGIs and minimally roam inactive domains (Højfeldt et al., 2019).

Interestingly, mESC differentiation is impaired in Pcl2 mutants and enhanced in Pcl3 mutants, leading to a proposal that different Pcl proteins may have different specificities for groups of target genes despite sharing many sites (Hunkapiller et al., 2012; Walker et al., 2010).

Our model raises the possibility that mouse Pcl proteins differentially control silencing by regulating PRC2 sampling rate. Simplistically, a Pcl with a low general affinity for CGIs could rapidly sample regulatory regions and promote silencing at lowly expressed genes similarly to core PRC2 in flies. In contrast, a Pcl with a higher general affinity for CGIs might more slowly sample them, antagonizing silencing and promoting the bivalent chromatin state observed in populations of progenitor cells. Although this simple model is certain to be further complicated by the relative abundances of different Pcl proteins and other subunits and the affinities of these subunits for every site in the genome, it provides a new framework to study how PRC2-dependent silencing could be modified by changes to PRC2 sampling rate induced by development or disease.

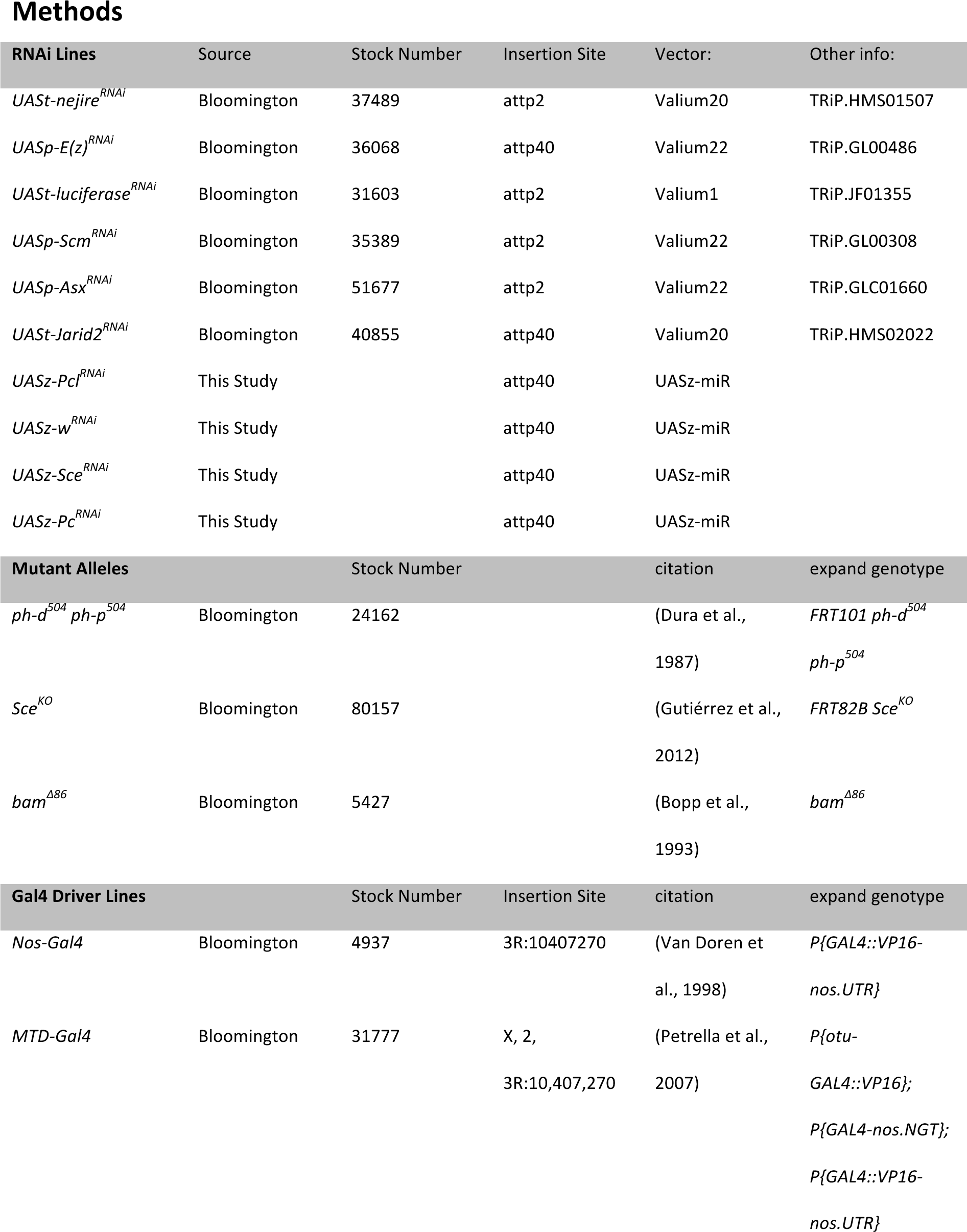
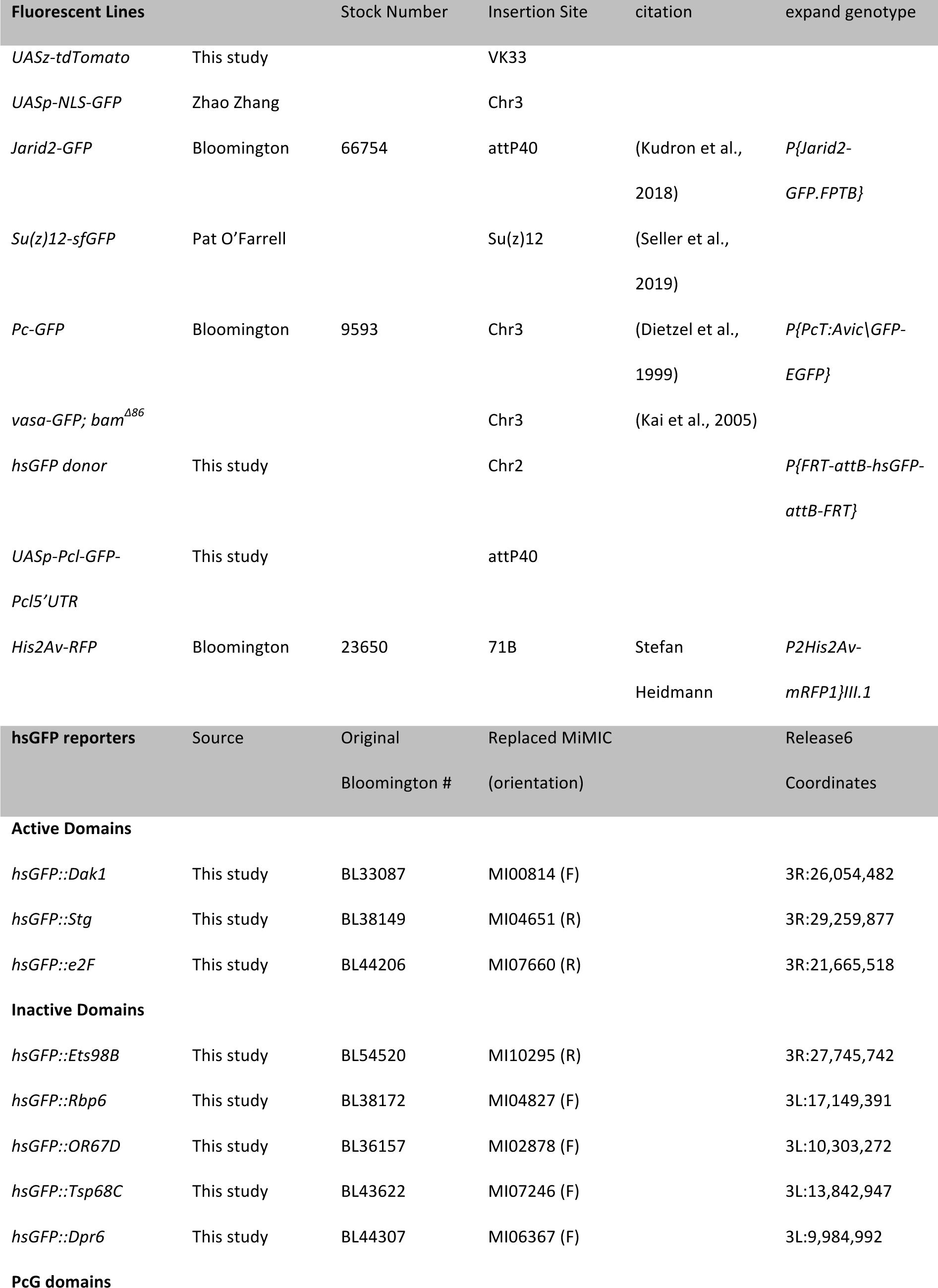
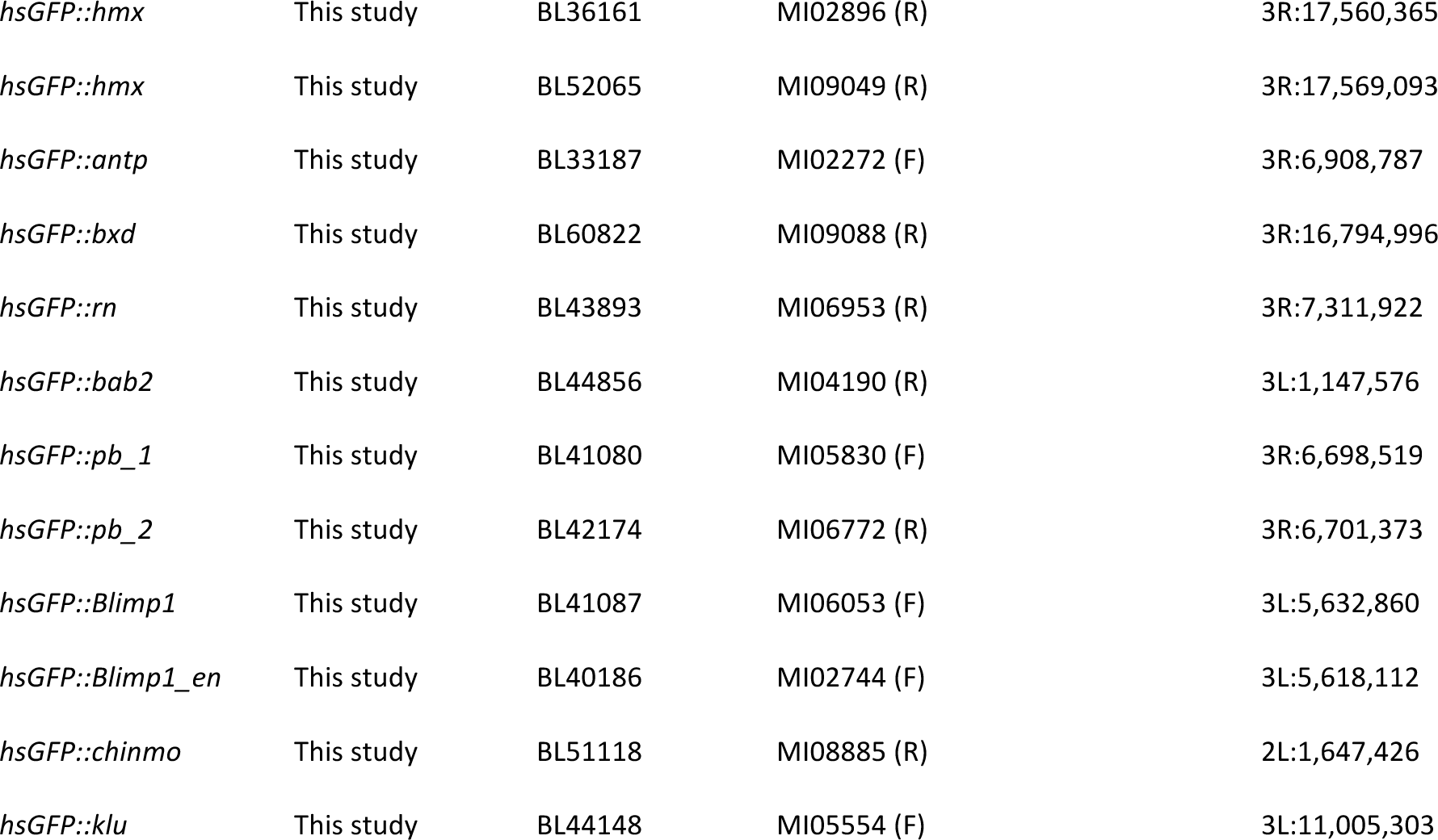

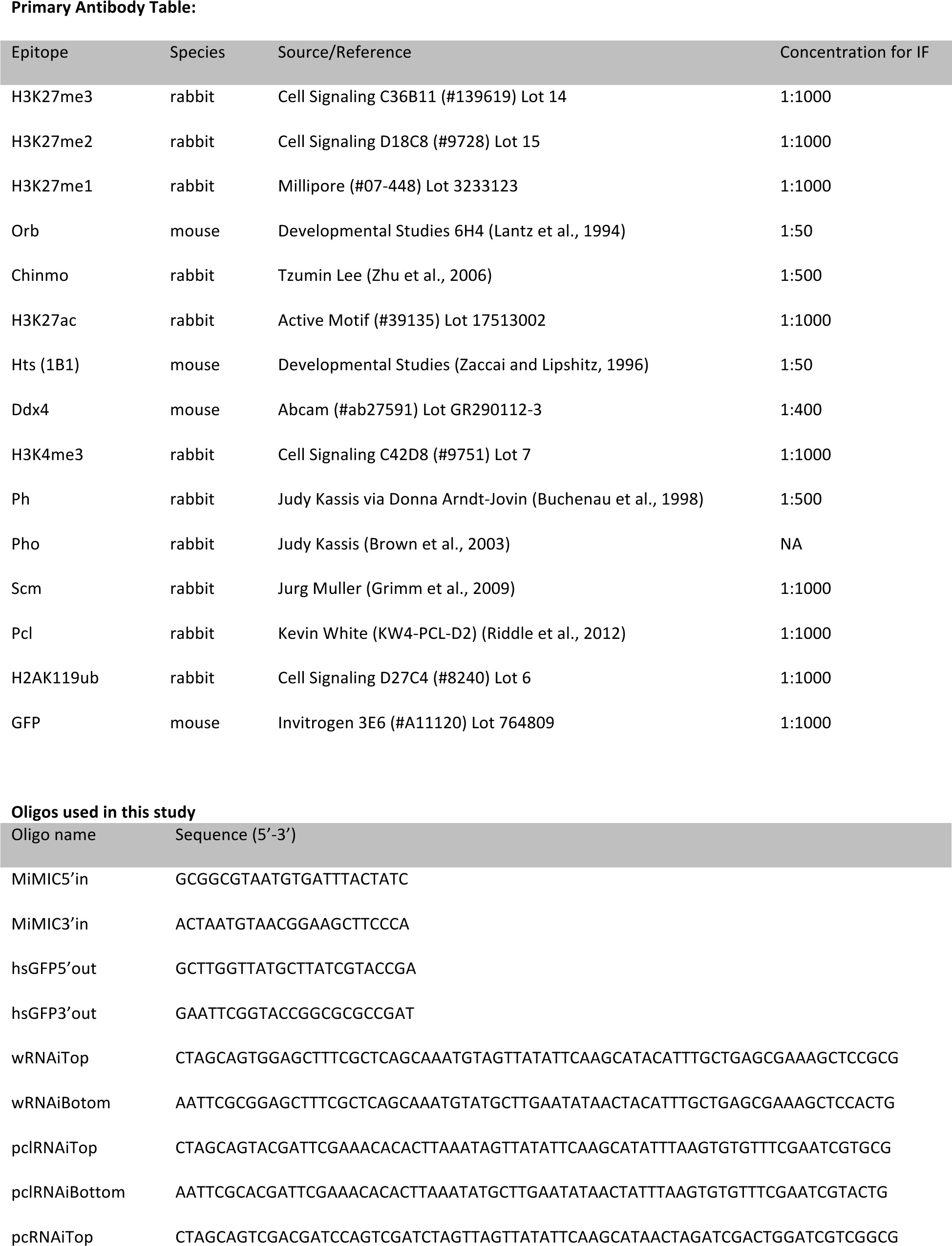
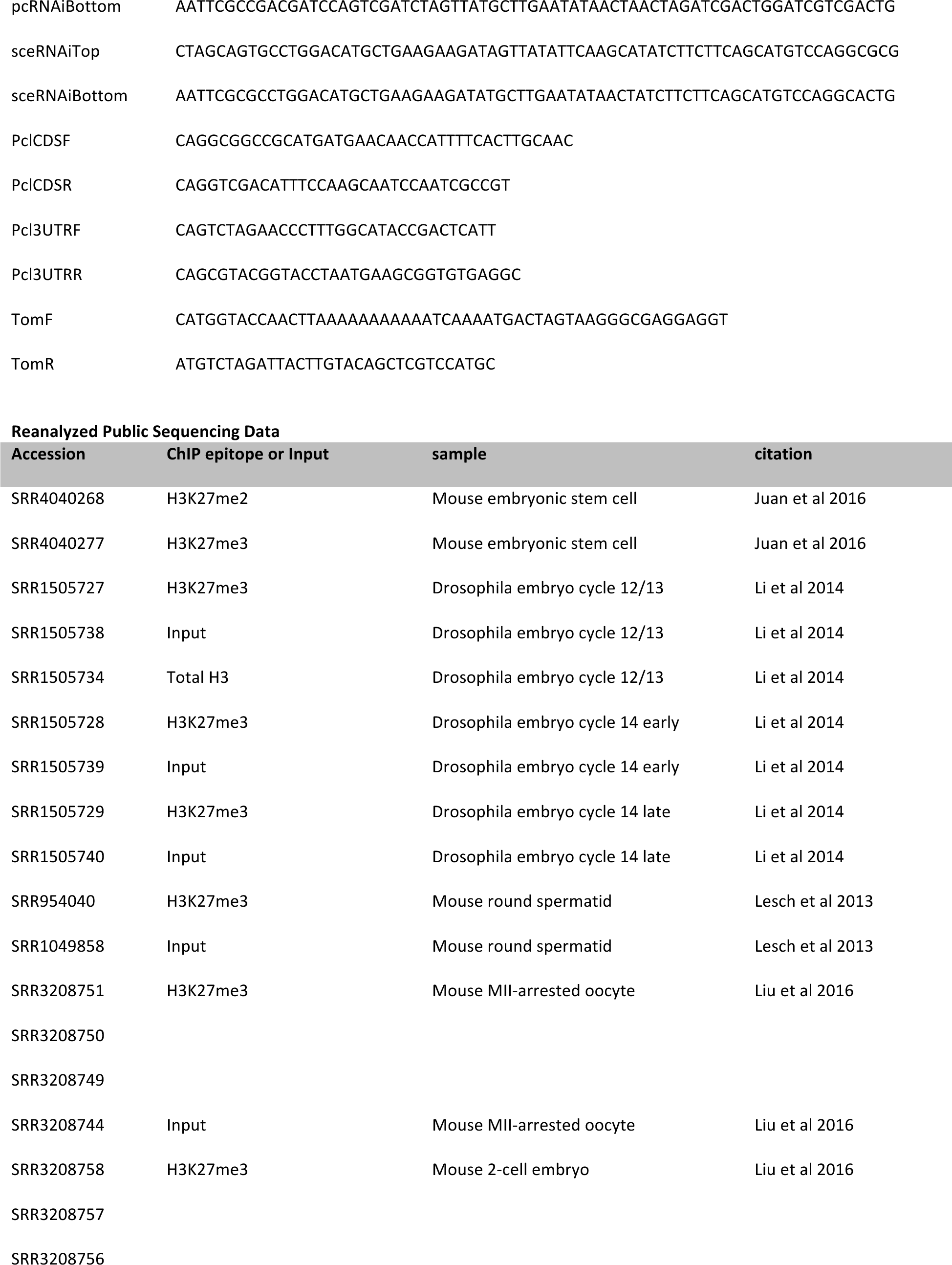
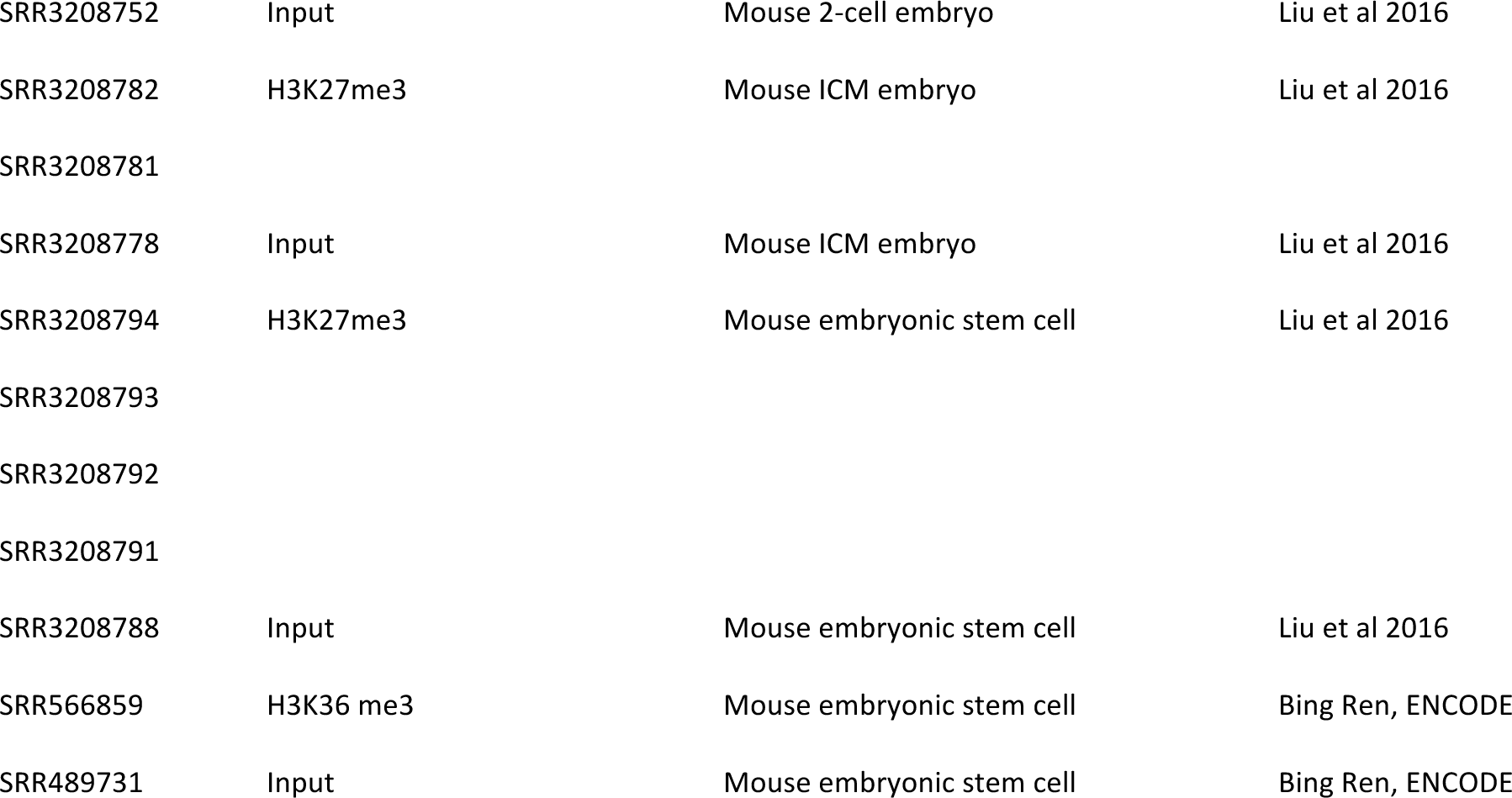

### Transgenic fly construction

To construct UASp-Pcl-GFP-Pcl3’UTR, we amplified Pcl coding sequence and introns from *OreR* (wild type) genomic DNA using PclCDSF + PclCDSR and cloned it between the NotI and SalI sites of pUASp-GFP-attB (DeLuca and Spradling 2018) to create pUASp-Pcl-GFP-attB. We then amplified the Pcl 3’UTR from genomic DNA using Pcl3UTRF + Pcl3UTRR and cloned it between the XbaI and BsiWI sites of pUASp-Pcl-GFP-attB to create pUASp-Pcl-GFP-Pcl3’UTR-attB. To construct pUASz-tdTomato-attB, we amplified tdTomato from pDEST-HemmarR (Han et al., 2011) with TomF + TomR and cloned it between the Acc65I and XbaI sites in pUASz-GFP-attB. To construct UASz-RNAi lines, we annealed Top and Bottom oligos and ligated the product between the NheI and EcoRI sites of pUASz-MiR. We introduced pUASz-tdTomato-attB into VK33 (3L:6442676) and all other attB-containing transgenes into attP40 (2L:5108448) using Rainbow Transgenics or BestGene Inc.

### Construction of hsGFP donor and introduction into existing MiMIC sites

GeneScript synthesized the hsGFP donor vector and we attached the sequence as a supplemental text file. hsGFP contains 400bp upstream and 40bp downstream of the Hsp70A transcription start site fused to a myosin intron, a synthetic translation enhancer, green fluorescent protein, and a P10 transcriptional terminator. We verified that both low and high-expressing hsGFP inserts were insensitive to *hsp70-*derived piRNAs by test crosses to *hsp70Δ* as in (DeLuca and Spradling, 2018). The hsGFP reporter is flanked on both ends by tandem attB (for phiC31-mediated recombination with MiMICs) and FRT (for mobilizing the donor construct from its initial locus) sites. To create initial donor lines on the second and third chromosome, we introduced hsGFP randomly into the genome through its truncated P-element terminal repeats (100bp 5’ and 195bp 3’) and P-element transposase-encoded helper plasmid. We isolated positive transformants by identifying GFP fluorescing flies after heat shock. To replace a MiMIC with our hsGFP reporter, we crossed a stable line carrying the hsGFP donor to flies carrying hsFLP, vasa-phiC31 integrase and appropriate marked chromosomes to generate F1 females carrying hsFLP, vasa-phiC31 integrase, the hsGFP donor, and appropriate marked chromosomes. We then crossed these F1 females to males carrying a yellow-marked MiMIC line of interest, and we heat-shocked the F2 eggs, larvae, and pupae at 37°C for 30 minutes every two days until adults eclosed. We then crossed F2 males carrying hsFLP, vasa-phiC31 integrase, the hsGFP donor, the yellow marked MiMIC recipient and an opposing marked chromosome, to yellow mutant females. We screened F3 progeny for flies that carry the original MiMIC chromosome but are yellow in color (i.e. MiMIC insertions where *yellow* was replaced by *hsGFP*) and generated stable lines. Generally, we obtained at least one independent replacement for every 5 F2 males undergoing MiMIC replacement.

Interestingly, the replacement success rate did not vary between MiMICs localized in different types of repressive chromatin, suggesting that germ cell precursors lack chromatin barriers to transgene insertion. We verified correct MiMIC replacement and determined the orientation of the reporter by PCR with primers flanking the hybrid attB/attP sites created by phiC31 recombinase.

### FACS sorting of live GSCs

We performed FACS sorting of live germline stem cells according to (Lim et al., 2012). Briefly, we dissected *Vasa-GFP; bam^Δ86^* ovaries in Grace’s Media + 10% fetal bovine serum and rinsed 2x in phosphate buffered saline (PBS). We dissociated cells by incubating with 0.5% Trypsin and 2.5 mg/ml collagenase for 13 min at room temperature with intermittent vigorous shaking. We washed 2x with PBS and twice filtered out large debris through a 50µm nylon mesh filter. We resuspended cells in PBS + 1% BSA + 1mM EDTA + 2ng/µl propidium iodide and sorted GFP positive, PI negative cells on a BD FACSAria III running FACSDiva software. After sorting, we spun down cells and prepared total RNA using the Ambion mirVana miRNA isolation kit (#AM1560) according to the manufacturer’s specifications without the miRNA enrichment step.

### FACS sorting nuclei for ChIP

We adapted a protocol from (Lilly and Spradling, 1996) to sort fixed, rather than live nuclei. We crossed *MTD-Gal4, UAS_z_-tdTomato* flies to *UAS_z_-w^RNAi^* (control) or *UAS-PcG^RNAi^* flies to generate F1 progeny heterozygous for the *MTD-Gal4* drivers, *UAS_z_-tdTomato*, and *UAS-RNAi*. To collect GSC-like progenitor nuclei, we crossed *nanos-gal4, UAS_p_-NLS-GFP, bam^Δ32^* heterozygous flies to control *bam^Δ32^* heterozygous flies or *UAS_z_-Pcl^RNAi^*, *bam^Δ32^* heterozygous flies to generate F1 females homozygous for *bam^Δ32^* and heterozygous for *nanos-gal4, UAS_p_-NLS-GFP,* and for experimental samples, *UAS_z_-Pcl^RNAi^*. We fed 3-7 day-old adult F1 females wet yeast paste for 3 days in the presence of males and dissected batches of 30-60 ovaries in ice-cold PBS. We treated each batch with 5mg/ml collagenase for 10 minutes at room temperature, and pipetted ovaries up and down to break up follicles and germaria. We rinsed 1x in PBS before fixing for 10 minutes in PBS + 2% paraformaldehyde at room temperature. We quenched fixation by adding 125 mM glycine for 5 minutes at room temperature, then washed in PBS. For *bam* ovary samples, we removed excess PBS, froze in liquid nitrogen, and stored at −80°C until we accumulated enough batches for our experiment. Before similarly freezing whole ovary samples, we passed the samples through a nylon mesh basket strainer to remove stage 10-14 follicles- which we found to interfere with subsequent fractionation steps. Once we accumulated enough batches, we thawed and resuspended samples in nuclear isolation buffer (NIB) + 0.1 µg/ml DAPI: 15 mM TrisHCl 7.4, 60 mM KCl, 15 mM NaCl, 250 mM sucrose, 1 mM EDTA, 0.1mM EGTA, 0.15 mM spermine, 0.5 mM spermidine, 1.5% NP40, protease inhibitor cocktail (Roche). We disrupted cells with 30 strokes in a dounce homogenizer with the B pestle, placed the extract on top of a 1M/2M sucrose step gradient, and spun at 20,000 x G for 20 minutes. After removing the supernatant, we resuspended the nuclear pellet in NIB with 20-40 more strokes of the dounce homogenizer with the B pestle (more strokes to dissociate smaller nuclei). We monitored nuclei dissociation after intermittent douncing by visualizing DAPI under a fluorescence microscope. We passed nuclei through a 100 µm filter, and diluted with 2 volumes of PBS before sorting nuclei in on a BD FACSAria III machine with 100 µm nozzle.

### Chromatin immunoprecipitation

For each IP, we started with a constant input of 0.5 million 2C-equivalent nuclei (for example, 0.5 million 2C cells or 2000 256C cells). We then added a spike in of 2,000-10,000 similarly fixed mouse 3T3 tissue culture cells or FACS-isolated *Drosophila pseudoobscura* 2-16C follicle cells. We resuspended nuclei in 100µl of 50 mM TrisHCl pH 8.0, 10 mM EDTA, 1% SDS, proteinase inhibitor cocktail (Roche) and sonicated in a Bioruptor Pico instrument with 22 cycles of 30 sec on, 30 sec off to fragment chromatin into mostly single nucleosome sizes (later confirmed with bioanalyzer). We then added 900µl of dilution buffer: 15 mM TrisHCl pH 8.0, 1 mM EDTA, 1% Triton X-100, 150 mM NaCl and saved 1% of this extract as input. For each IP, we preincubated 10µl of antibody with 25µl of a 1:1 mix of proteinA:proteinG dynabeads and washed 2x with PBS + 0.02% Tween 20 (PBST). We combined antibody-conjugated beads with chromatin extracts on a rocker at 4°C overnight. Washed 2x 15 min each with **Wash buffer A**: 20 mM TrisHCl pH8.0, 2 mM EDTA, 0.1% SDS, 1% Triton X100, 150 mM NaCl, **Wash buffer B:** 20 mM TrisHCl pH8.0, 2 mM EDTA, 0.1% SDS, 1% Triton X100, 500 mM NaCl, **Wash buffer C:**10 mM TrisHCl pH8.0, 1 mM EDTA, 1% NP40, 1% Sodium deoxycholate, 0.25M LiCl, **TE buffer:** 10 mM TrisHCl pH8.0, 1 mM EDTA. We eluted chromatin and reversed crosslinking by incubating at 65°C overnight with **Direct Elution Buffer (DEB):** 10 mM TrisHCl pH 8.0, 300 mM NaCl, 5 mM EDTA, 0.5% SDS. We additionally reversed crosslinks in input samples by adding NaCl to 300mM and adding DEB to equalize the volume between inputs and IPs and incubating at 65°C overnight. We then treated samples for 30 min at 37°C with 0.3mg/ml RNAse A, and 2 hours at 55°C with 0.6 mg/ml proteinase K, before extracting DNA with phenol:chloroform and precipitating with NaAc/ethanol. After a 70% ethanol wash, we resuspended samples in 10µl water and used all 10µl for library prep.

### RNA preparation for RNAseq

We crossed MTD-Gal4 females to UAS-RNAi males to generate F1 females heterozygous for the MTD-Gal4 drivers and a UAS-RNAi transgene. Because *E(z)^GLKD^* ovaries begin to degenerate at stage 6, we dissected ovaries from control *Luciferase^GLKD^* or *E(z)^GLKD^* females 0-8hrs after they eclosed from the pupal case. At this time point, both *control* and *E(z)^GLKD^* follicles had not progressed past stage 6 and therefore contained a nearly identical distribution of stages and cell types. The other *PcG^GLKD^* ovaries analyzed in this paper did not degenerate at a particular stage, so we compared RNA from fully developed ovaries in control *w^GLKD^* and *PcG^GLKD^*. We fed 3-7 day old females fed wet yeast paste for 3 days before dissecting ovaries in cold PBS. We dissected 30 ovaries per replicate for ovaries containing nurse cell stages and 50 ovaries per replicate for *bam* mutant ovaries. We purified total RNA using the TriPure reagent (Sigma Aldrich) according to the manufacturer’s protocol.

### Library preparation and sequencing

For RNAseq, we used the Illumina TRUseq version 2 kit to create polyA enriched mRNA libraries according to the manufacturer’s specifications. For ChIPseq, we used the Takara Bio ThurPLEX DNA seq kit according to the manufacturer’s specifications.

Briefly, double stranded DNA ends are repaired, universal adapters are ligated on, and indexing is performed in a single tube using a total number of 15 PCR cycles. We sequenced both ChIP and RNAseq libraries on an Illumina NextSeq 500 using 75bp single end reads.

### Live cell imaging of Pcl-GFP

We collected embryos from *Nanos-Gal4, His2A-mRFP/+, UASp-PclGFP-Pcl5’UTR/+* flies for 1 hour on standard apple juice agar plates and aged for 30 minutes before dechorionating with 50% bleach for 2 min in a basket strainer and rinsing with water until we did not detect bleach smell. We attached embryos to a #1.5 coverslip with homemade glue made by dissolving double-stick tape in heptane and coated embryos in gas-permeable halocarbon oil. For each experiment we simultaneously imaged GFP and RFP in 10-15 embryos every 3-5 minutes on an inverted spinning disc confocal with 20x Plan Apo 0.8 NA objective and dual EMCCD detectors.

### Ovary fixation for antibody staining and in situ hybridization

Before ovary fixation, we fed 3-7 day-old adult females wet yeast paste for 3 days in the presence of males. We dissected ovaries in PBS and fixed in PBS + 4%paraformaldehyde + 0.01% TritonX-100 for 13 minutes before washing 2x with PBS + 0.1% TritonX-100 (PTX).

### Antibody staining

We blocked fixed ovaries with PTX + 5% Normal Goat Serum (NGS) for 30 min. We incubated ovaries with primary antibody at 4°C overnight in PTX + 5% NGS, washed 3x in PTX, then incubated with alexa-fluor conjugated secondary antibody (1:1000) in NGS at 4°C overnight. We washed 3x in PTX, including 0.5 µg/ml DAPI in the second wash, and mounted in 50% glycerol. A list of primary antibodies and concentrations is listed in Table 1.

### In Situ Hybridization

We ordered custom Stellaris RNA FISH oligonucleotide probes directed against Pcl mRNA conjugated to CALFluor RED 590 from LGC Biosearch Technologies and performed *in situ* hybridization based on the company’s recommendations. Briefly, we washed fixed ovaries 2x 20 min. in Wash Buffer A (LGC Biosearch Technologies #SMF-WA1-60) + 10% formamide (WAF) then incubated ovaries for at least 2 hours in hybridization buffer (LGC Biosearch Technologies #SMF-HB1-10) + 10% formamide (HBF) at 37°C. We then incubated ovaries in HBF + 50 nM probe overnight, before washing 1x with HBF, 3x WAF at 37°, and 2x Wash Buffer B (LGC Biosearch Technologies #SMF-WB1-20) at room temperature. We mounted ovaries in 50% glycerol and imaged on a Leica SP8 scanning confocal.

### hsGFP Reporter Assay

We recombined each hsGFP reporter insertion with nosGal4 using standard fly genetics. For each experiment, we crossed hsGFP+NosGal4 females to UAS-RNAi males and assayed reporter induction in F1 progeny carrying the hsGFP, NosGal4, and UAS-RNAi. We cultured flies at 22°C to prevent premature activation of the heat shock response and allow *E(z)^GLKD^* ovaries to produce rare follicles progressing past stage 6. For each experiment, we collected 3-7 day-old females and fed them wet yeast paste in the presence of males for 3 days to achieve a maximum rate of egg production. For each line, we heat shocked half of the well-fed flies while using the other half as a no heat shock control. We heat shocked flies in vials containing 1mL of solidified 1% agar in a 37°C water bath for 20 min. After heat shock, we returned flies to normal food plus wet yeast for 3 hours to allow for GFP protein production and maturation. We dissected whole ovaries in PBS and fixed for 30 min in PBS + 4% paraformaldehyde + 0.01%Triton X-100. After 1 wash in PBS+0.1%TritonX-100 (PTX), we treated ovaries in PTX + 100µg/ml RNAse A for 1-2 hours at room temperature before staining with 0.2µg/ml Propidium Iodide to visualize DNA for developmental staging. We mounted ovaries in 50% glycerol and directly imaged GFP and Propidium Iodide fluorescence on a spinning disc confocal on a spinning disc confocal with 20x Plan Apo 0.8 NA objective. We acquired a z-stack of confocal images and chose a single confocal plane through the middle of the desired germline stem cell or cyst and measured mean GFP fluorescence in germ cells in a manually drawn region of interest in image J. For each line and condition (+/-heat shock, different RNAi), we imaged at least 3 independent ovary pairs under identical laser power and acquisition settings and measured mean GFP fluorescence intensity in 1-5 germline stem cells or germ cell cysts of each stage per ovary pair. For each reporter line, to calculate a single experimental replicate for “induction at a given stage,” we first determined the mean intensity of all replicates of the non-heat shocked measurements for that stage and subtracted this mean from each experimental replicate for that stage.

### Genome Segmentation

We converted a bed file of the 9 state chromatin model for S2 cells from (Kharchenko et al., 2011) to Drosophila genome release 6.02 coordinates and simplified the 9 chromatin state model into a 4 state model containing an active compartment (States 1-5) a PcG compartment enriched for PcG proteins and H3K27me3 (State 6), an inactive compartment enriched for H3K27me2 (State 9), and a Hp1 compartment enriched for Hp1a, Su(var)3-9, and H3K27me2/3 (State 7-8).

To annotate genes and transcription start sites, we used BEDTools (Quinlan and Hall, 2010) to intersect the 4 state model with the *Drosophila* genome release 6 annotation from Ensembl 81 to append a chromatin domain type to each protein-coding gene or transcription start site (TSS), removing any gene or TSS residing on the Y-chromosome, the 4^th^ chromosome, pericentric heterochromatin on any arm. Because genes have multiple isoforms that may span multiple domain types, we simplified the coordinates of the many isoforms of each gene into a single coordinate spanning the largest region shared by all isoforms. We then intersected these coordinates with the coordinates of PcG domains, retaining any gene with any bit of PcG domain as a PcG gene. We intersected the remaining genes with the coordinates of active domains, retaining any gene with any bit of an active domain as an active gene. We classified the remaining genes as inactive genes.

For ChIP analysis, we segmented the genome into 5 kilobase bins that start every 500 bases. We removed bins from the Y chromosome, the 4^th^ chromosome, or any scaffolds not part of the remaining 5 chromosome arms. We additionally removed bins residing in 3 most heavily amplified chorion gene clusters. We annotated the remaining bins with a single chromatin type by intersecting the list of bins with our 4-state model, excluding any bin that contained multiple chromatin types from further analysis. Thus, the edges of chromatin domains were slightly underrepresented in our bin analyses. In Figure 6, we further classified PcG bins to separate bins that also contained a PRE. We classified PRE bins as any PcG bin intersecting with a composite list of Pho plus Ph peak summits called by MACS2.

### RNAseq Analysis

We aligned 75 base pair single-end reads from at least 3 control and 3 experimental replicates to the Drosophila release 6 genome, Ensemble 81 annotation using the default parameters of Hisat2 (Kim et al., 2015). We measured raw transcripts per million for each gene using the default parameters of StringTie (Pertea et al., 2015). For differential expression analysis, we extracted raw read counts mapping to each gene, generated a DEseq2 model (Love et al., 2014), and analyzed and plotted normalized abundances of protein coding genes in R.

### ChIPseq Analysis

We used bowtie2 (Langmead and Salzberg, 2012) to align 75 base pair single-end reads to either the *Drosophila melanogaster* release 6 genome (when no spike in was present) or a hybrid genome of *Drosophila melanogaster* and *Drosophila pseudoobscura* or *Mus musculus* (when corresponding spike in was present). We used the proportion of spike-in reads in Input and IP samples to generate a normalization factor for subsequent ChIPseq analysis and used a MAPQ 30 filter to remove ambiguously mapped reads. After scaling read coverage to reads per million and applying the normalization factor, we visualized and presented read depth across a genomic region of interest in the Integrative Genomics Viewer (Robinson et al., 2011), or generated genome wide summaries as described below.

To calculate read density across different chromatin domains, we used bedtools to assign spike-in normalized read coverage in Input and IP samples to annotated overlapping 5kb bins. We used ggplot2 to plot a smoothed histogram of enrichments (IP/In) across all bins, with the y-axis corresponding to the proportion (not raw number) of bins for a given domain type. We used MACS2 (Zhang et al., 2008) to call peaks in Pho and Ph IP samples and Deeptools (Ramírez et al., 2014) to present raw read depth heat maps of genomic regions surrounding all peaks.

## Acknowledgements

We thank Judy Kassis for Pho and Ph antibodies, Jurg Muller for Scm antibody, and Kevin White for Pcl antibody. We thank Patrick O’Farrell for Su(z)12-sfGFP, Zhao Zhang for UASp-NLS-GFP, and the Bloomington Stock Center and the Drosophila Gene Disruption project for other fly stocks. We thank Allison Pinder, Frederick Tan, and John Urban for help in generating and analyzing high throughput sequencing data. S.Z.D was supported by a Helen Hay Whitney Fellowship and M.G. was supported by a Jane Coffin Childs Fellowship. Correspondence and requests should be addressed to A.C.S.

## Supplementary Files

**Fig S1:**
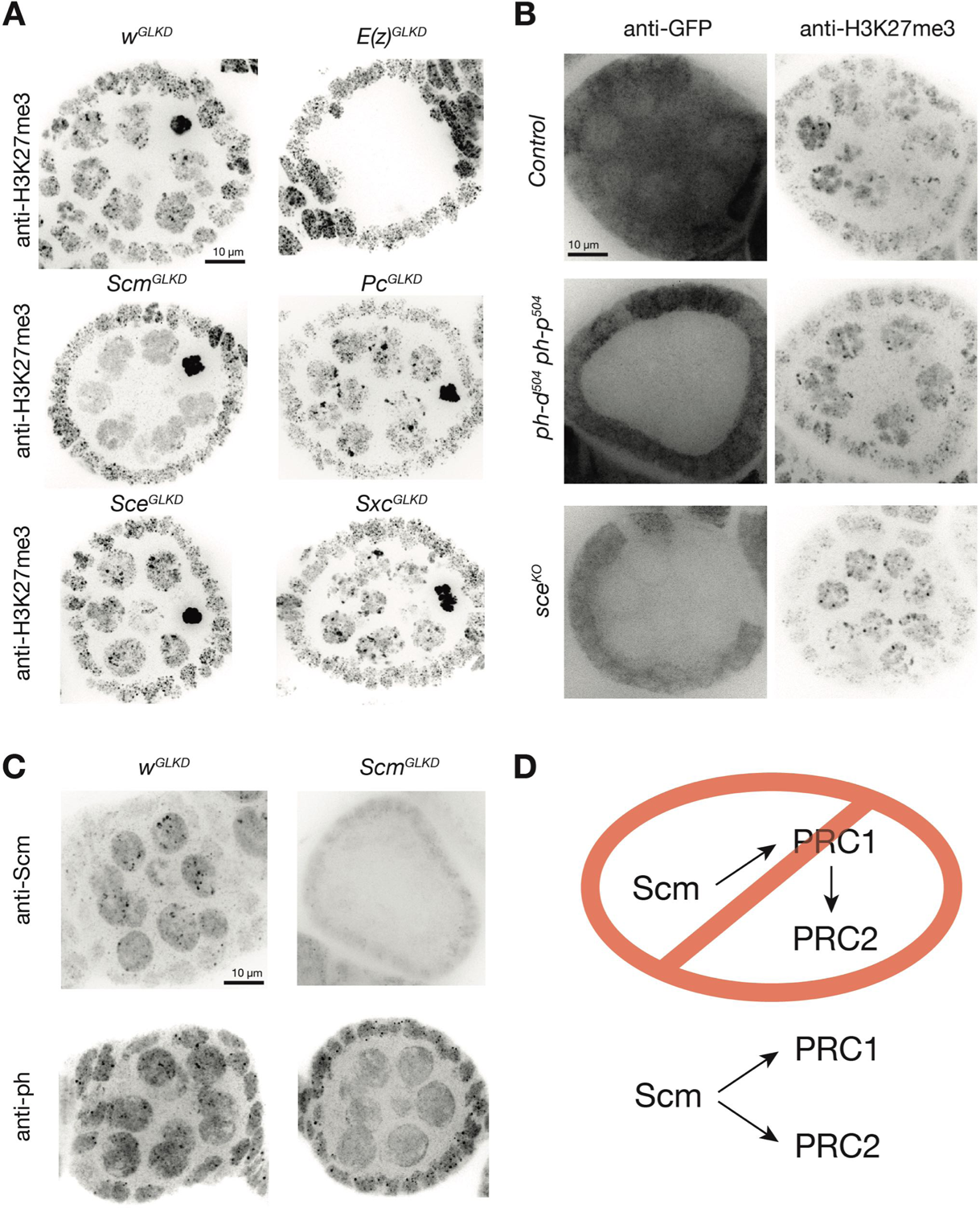
**Scm promotes H3K27me3 enrichment independently of PRC1** (A) H3K27me3 staining of stage 4/5 follicles showing punctate pattern of H3K27me3 staining in negative control (*w^GLKD^*) or PRC1-depleted (*Pc^GLKD^*, *Sce^GLKD^*) nurse cells. Pronounced puncta were absent from *Scm^GLKD^* nurse cells or positive control *E(z)^GLKD^* nurse cells. (B) *ph-d/ph-p* or *sce* null mutant clones were generated by mitotic recombination and visualized by lack of GFP fluorescence. H3K27me3 foci were unaltered by complete removal of Ph or Sce activity. (C) IF images showing *Scm^GLKD^* effectively removes Scm protein from nurse cells and prevents the coalescence of Ph into discreet foci. (D) Model for how PRC2 is targeted to PcG domains by Scm independently of PRC1.

**Fig S2.**
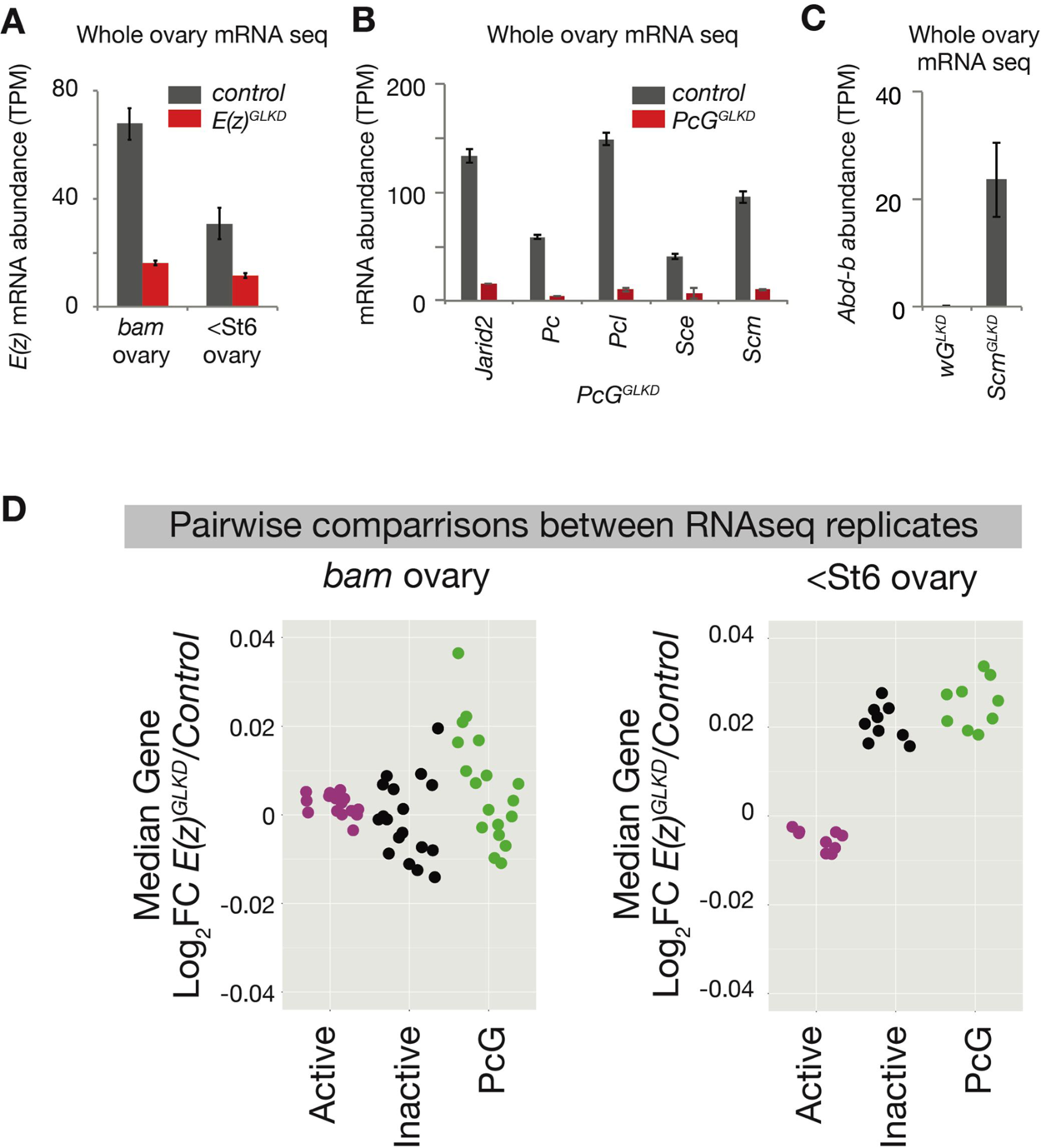
RNA seq controls (A) RNAseq-measured *E(z)* mRNA abundance in *Luc^GLKD^* (control) or *E(z)^GLKD^* whole ovaries. Note that residual *E(z)* mRNA in *E(z)^GLKD^* may originate from somatic cells surrounding germline tissue in both *bam* mutant and *bam+* ovaries containing follicles up to and including stage 6. (B) RNAseq-measured mRNA abundance of the indicated *PcG* genes in fully developed whole ovaries (up to and including stage 14) following GKLD of control (*w*) or the indicated *PcG* gene. Note that knockdowns in (B) appear to be more efficient than those in (A) because the ratio of germline to soma cytoplasmic volume increases as follicles grow. (C) RNAseq-measured abundance of *Abd-b* in control *w^GLKD^* or *Scm^GLKD^* whole ovaries. (D) Pairwise comparison of the median fold change in active, inactive, and PcG domain genes between individual experimental replicates.

**Fig S3.**
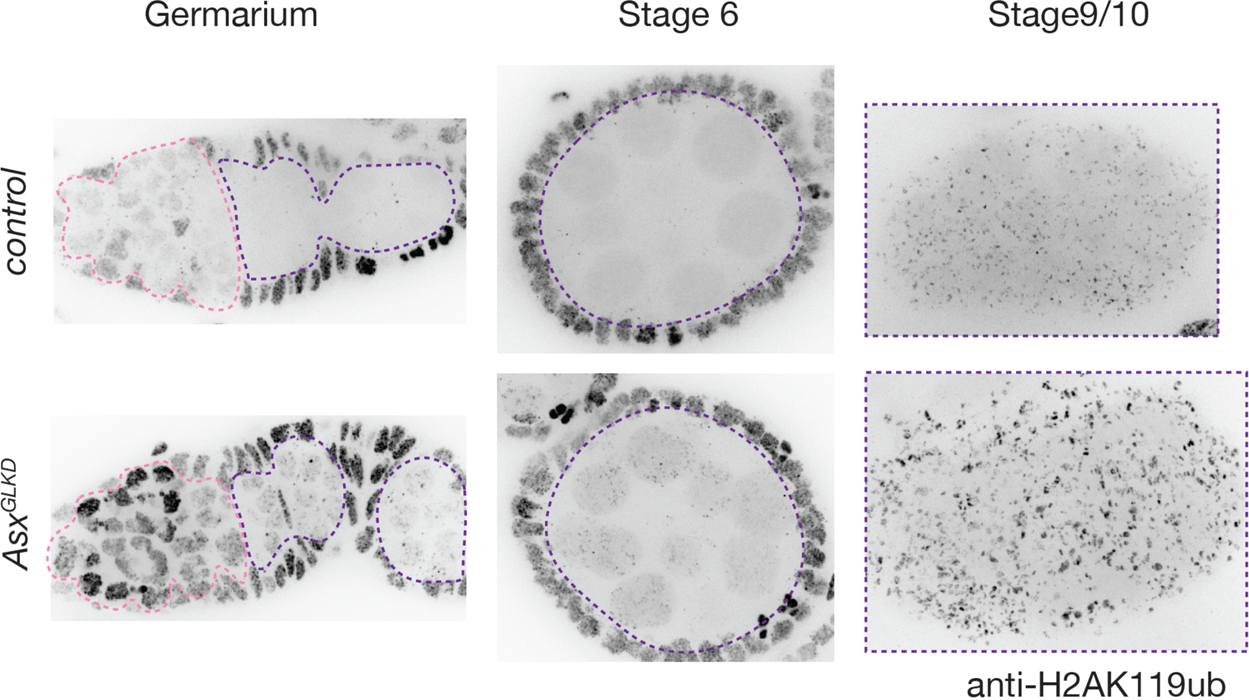
H2Ak119ub levels during nurse cell development Immunofluorescent images of H2Ak119ub distribution in control (*w^GLKD^*) or *Asx^GLKD^* ovaries.

**Fig. S4.**
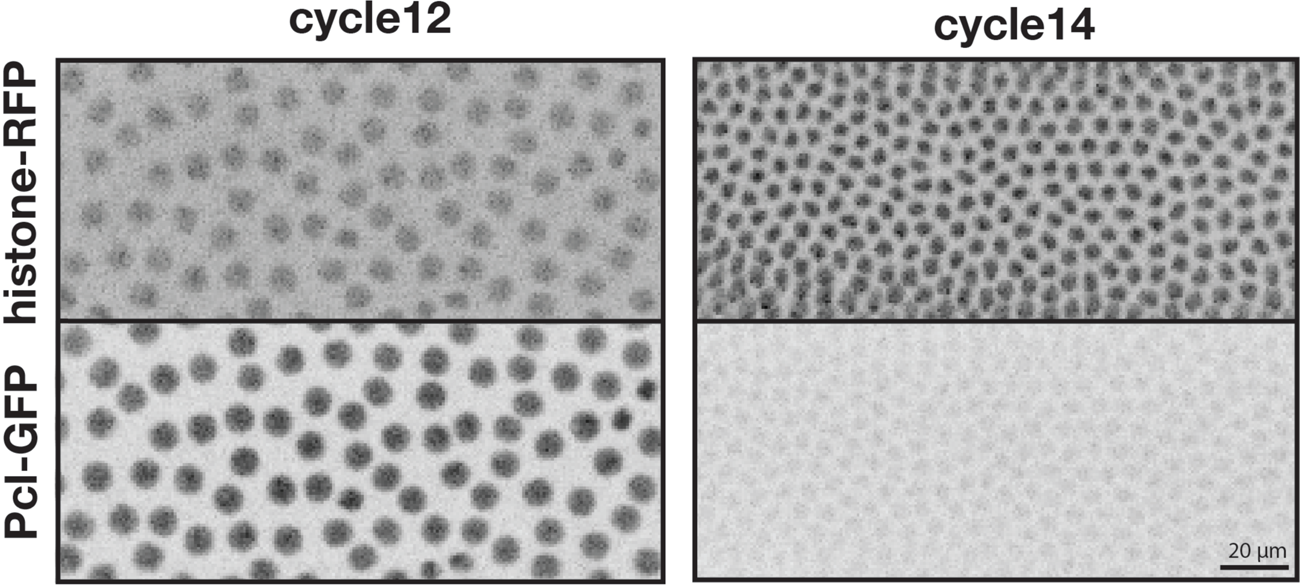
Pcl depletion during early embryogenesis Live embryos containing histone-RFP to stage the cell cycle and maternally inherited Pcl-GFP from *Mtd>Gal4 UASp-PclGFP-Pcl5’UTR* mothers. Pcl protein is abundant in nuclei before the maternal to zygotic transition at cycle 14 but promptly decreases at cycle14.

**Fig. S5.**
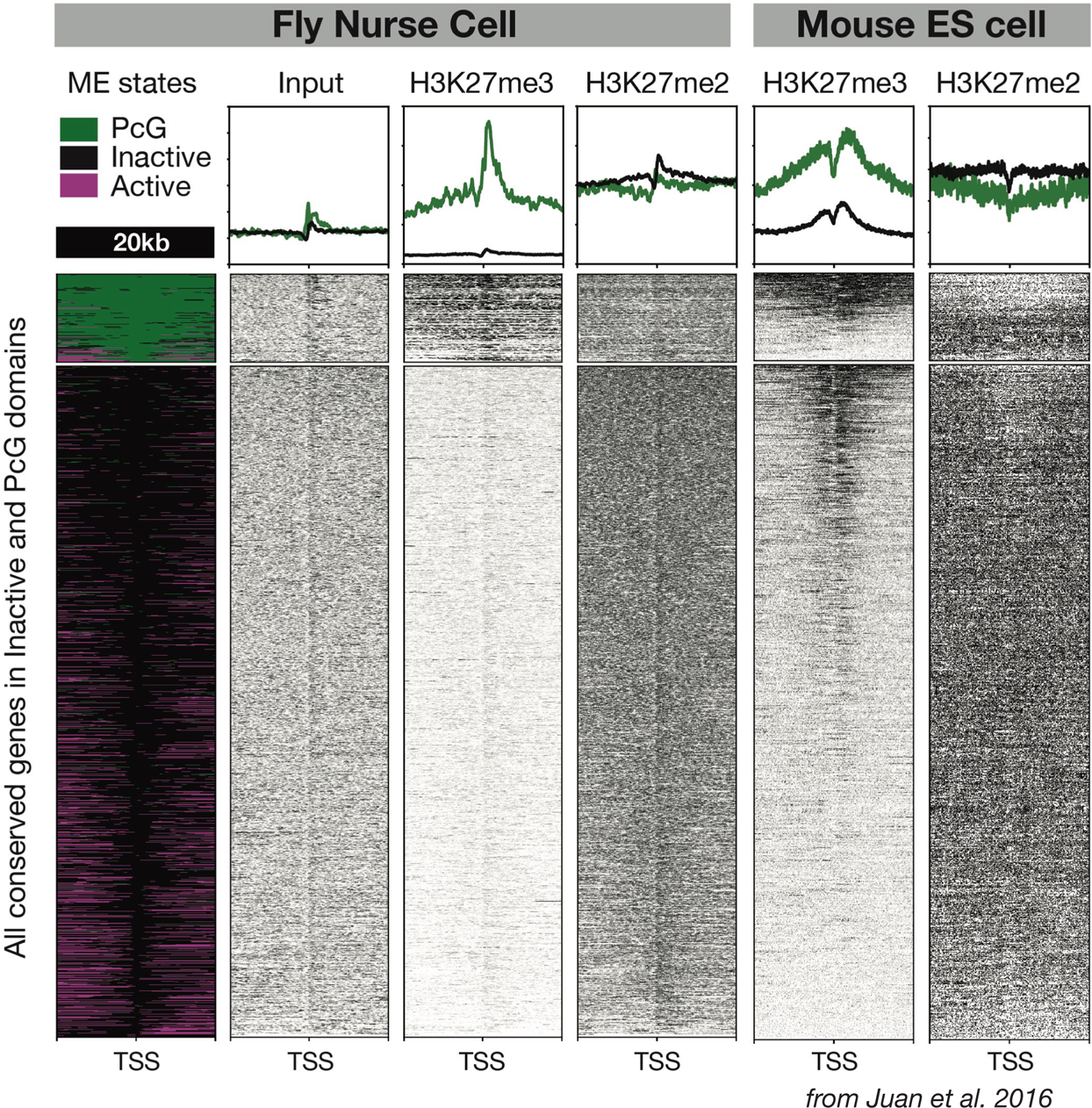
H3K27 methylation patterns across homologous genes in flies and mice The left four columns plot 1652 fly genes residing in PcG or Inactive domains while the right two columns plot the direct mouse homologues of those genes using data from (Juan et al., 2016). Each row is a 20kb genomic region centered on the transcription start site of a gene (TSS) with its coding region to the right. Genes in PcG domains (green) are independently arranged from genes in inactive domains and the mean signal across each domain type is plotted above the raw read depth heatmaps. Note that fly TSSs in inactive domains have minor H3K27me3 enrichment while many of their mouse homologues have substantial H3K27me3 enrichment.

**Fig. S6.**
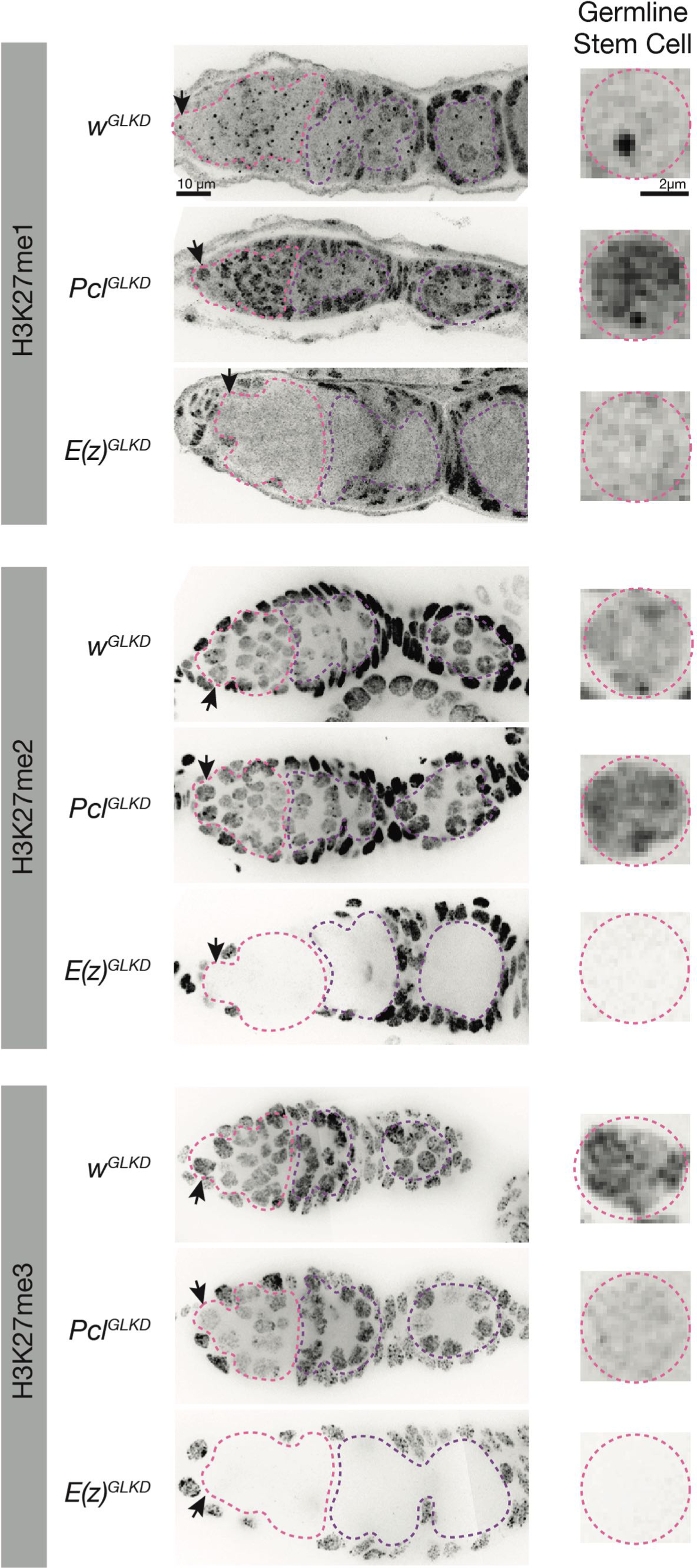
Pcl promotes higher H3K27 methylation states in germline precursors IF images of ovaries of the indicated genotype stained for the indicated H3K27me epitope. Germline progenitors are outlined in pink and nurse cells are outlined in purple. Arrows point to a single germline stem cell nucleus for each condition that is magnified on the right.

